# *Vibrio* spp. dominate the microbiome of the endosymbiotic algae in healthy coral tissues

**DOI:** 10.1101/2025.08.07.668945

**Authors:** Abigail Smith, Kendall N. Jung, Catherine Keim, Lea G. Stahr, Molly Schechter, Adejumoke Ewedemi, Anny Cárdenas

**Author notes:** Authors contributed equally (Abigail Smith, Kendall N. Jung).

## Abstract

Coral reefs are rapidly declining due to climate change, and natural recovery mechanisms can no longer keep pace. Coral probiotics have emerged as a promising restoration tool, yet their broad application is limited by our incomplete understanding of long-term microbial associations crucial to coral health. Microbes closely associated with the coral algal symbionts (Symbiodiniaceae) are particularly promising candidates, given their potential to enhance algal function and host resilience. Although recent studies have begun to characterize these Symbiodiniaceae-associated microbial communities, methodological differences in algal cell purification have led to inconsistent results. Here, we compared multiple sample processing steps to generate Symbiodiniaceae-enriched fractions from clonal fragments of *Acropora nobilis*, and examined the resulting microbial communities. We consistently detected members of Flavobacteraceae, Rhodobacteraceae, Rhizobiaceae, and the genus *Marinobacter*. Strikingly, *Vibrio* species dominated the Symbiodiniaceae fractions across protocols. We isolated and sequenced 11 *Vibrio* strains enriched in these fractions and identified genes related to both virulence and putative beneficial traits, including vitamin biosynthesis and antioxidant production. Despite confirming their high potential for virulence, the persistent association, spatial proximity to Symbiodiniaceae, and presence of genes suggesting beneficial functions point to possible mutualistic roles for these *Vibrio* strains within the coral holobiont. This work highlights the value of fraction-based sampling for resolving microbiome structure, and emphasizes the need to reassess the ecological roles of key microbial taxa. By advancing our understanding of Symbiodiniaceae-bacteria interactions, this work provides a foundation for exploring microbial contributions to coral health and resilience.

## INTRODUCTION

Symbiodiniaceae are a diverse family of photosynthetic microalgae that live endosymbiotically with various marine organisms, but are best known for their symbiosis with corals (Kirk and Weis 2016; LaJeunesse et al. 2018; Goulet, Lucas, and Schizas 2019). This symbiosis is crucial for the success and diversity of corals but is also vulnerable to environmental stress (Muscatine, R. McCloskey, and E. Marian 1981). The loss of Symbiodiniaceae, known as bleaching, affects many marine organisms, and while the most common cause for it is ocean warming (Hughes et al. 2017; Rädecker et al. 2021), it can also be triggered by eutrophication, deoxygenation, and ocean acidification (Douglas 2003; Alderdice et al. 2022; Anthony et al. 2008). Coral bleaching is the primary cause of coral death worldwide, often resulting in the collapse of entire coral reef ecosystems (Stuart-Smith et al. 2018; Suggett and Smith 2020) with devastating environmental and social consequences (Berkelmans et al. 2004; Camp et al. 2018).

Symbiodiniaceae undoubtedly contribute to coral thermal susceptibility (Cunning and Baker 2012; Rädecker et al. 2021; Silverstein, Cunning, and Baker 2015), making it imperative to understand their diversity and functioning to elucidate bleaching processes and develop effective conservation strategies for coral reefs. In particular, identifying the biological aspects that distinguish thermally resistant from sensitive Symbiodiniaceae species is fundamental to supporting current coral conservation efforts (Santoro et al. 2025; Amario et al. 2023). However, some aspects of Symbiodiniaceae biology, particularly their interactions with other microbes in coral tissues, are still poorly understood (Garrido et al. 2021; Matthews et al. 2020). In this regard, most coral microbiome studies have used “bulk” approaches that typically do not consider the compartmentalization of coral tissues into discrete micro-habitats. These micro-habitats are spatially structured at finer scales and harbor specific and specialized bacterial communities (Pernice et al. 2020; Cárdenas et al. 2022; Pollock et al. 2018). For example, bacterial communities that reside inside the symbiosome, a specialized compartment within the coral gastrodermal cell that houses Symbiodiniaceae, are different from the communities within other parts of the coral and have only recently attracted scientific interest (Maire et al. 2021; Maire, Blackall, and van Oppen 2021; Garrido et al. 2021). Yet, much remains unknown about these bacteria and their interactions with the algae.

Current knowledge of Symbiodiniaceae-bacterial interactions is mainly derived from long-term cultures of Symbiodiniaceae (i.e., *ex hospite*) that have adapted to a free-living lifestyle compared to Symbiodiniaceae cells living in a host cell (i.e., *in hospite*). Bacterial communities associated with Symbiodiniaceae *ex hospite* are numerous, diverse, and arguably essential, given the difficulty of growing axenic (i.e., bacteria-free) cultures (Ritchie 2011; Lawson et al. 2018; Camp et al. 2020). While microbiomes of *ex hospite* Symbiodiniaceae are species-specific and likely differ from those *in hospite*, some bacterial groups are conserved across Symbiodiniaceae cultures and are also prevalent in coral tissues. For instance, members of the genera *Marinobacter,* are commonly shared across Symbiodiniaceae cultures (Lawson et al. 2018; Maire et al. 2023; Matthews, Khalil, et al. 2023) and also presumed to play roles in coral development given their high prevalence in larvae (Sharp, Distel, and Paul 2012) as well as in the skeletons of adults (Cárdenas et al. 2022). Similarly, *Labrenzia* has been identified as a common associate in Symbiodiniaceae cultures (Lawson et al. 2018; Buerger et al. 2022), but has also been isolated from coral tissues where it has been proposed to play a role in sulfur cycling (Kuek et al. 2022; Raj Sharma et al. 2019; Gardner, Leggat, and Ainsworth 2023). Interestingly, both bacterial groups have been found to produce the plant hormone indole-3-acetic acid (IAA), that triggers algal growth in Symbiodiniaceae-bacterial co-cultures (Matthews, Hoch, et al. 2023; Matthews, Khalil, et al. 2023).

Recent studies have provided the first insights into the *in hospite* microbiomes of Symbiodindiniaceae, revealing that members of the classes Flavobacteriia, Alphaproteobacteria, and Gammaproteobacteria, particularly those from the families Flavobacteriaceae, Rhodobacteraceae, and Alcanivoracaceae, as well as members of the order Rhizobiales, are commonly dominant within these spatially distinct bacterial communities (Maire et al. 2021; Hill et al. 2024; Gardner, Leggat, and Ainsworth 2023; Ainsworth et al. 2015). However, a comprehensive understanding of the diversity, stability, and functional roles of *in hospite* Symbiodiniaceae microbiomes, and how these microbial associates interact with their dinoflagellate hosts, remains limited. One of the main barriers to advancing this knowledge is the lack of a simple, reproducible, and scalable method for isolating *in hospite* Symbiodiniaceae and their closely associated microbiomes. Without a standardized approach, comparisons across studies are challenging, and key microbial taxa may be overlooked or inconsistently detected. To address this gap and improve our understanding of *in hospite* Symbiodiniaceae-associated microbiomes, we systematically compared the microbial communities recovered under different Symbiodiniaceae isolation protocols. Specifically, we evaluated how centrifugation strategies, sample processing conditions, and nucleic acid templates influence microbial composition. Our study provides the first direct comparison of these variables, offering a framework for optimizing the recovery of *in hospite* microbiomes. In doing so, we identify bacterial taxa that are consistently enriched across protocols and therefore most likely represent core members of the Symbiodiniaceae-associated microbiome.

## MATERIALS AND METHODS

### Sample fractionation protocols

Twelve clonal fragments of *Acropora nobilis* were purchased from the World Wide Corals aquaria facility in Florida. Upon arrival at the laboratory, six fragments were processed immediately, while the remaining six were snap-frozen in liquid nitrogen and stored at −80 °C for later processing. Each fragment was placed in a sterile plastic bag with 6 ml of phosphate-buffered saline (PBS) containing 2% NAC. Tissue was removed from the skeleton using air-blasting with compressed air delivered through sterile pipette tips, continuing until the skeleton was visually bleached. The resulting slurry was collected in 50 ml tubes, homogenized using a handheld tissue homogenizer (VWR) for 30 seconds, and centrifuged at 500 g for 30 seconds to remove large debris. Aliquots were collected at this stage to represent the unfractionated sample (F1). The remaining supernatant was evenly distributed into fresh 15 ml tubes for testing two centrifugation protocols that reflect commonly used approaches previously described (Maire et al. 2021; Gardner, Leggat, and Ainsworth 2023): one slow and long, the other fast and short. Protocol 1 (P1) involved centrifugation at 500 g for 5 minutes, while protocol 2 (P2) used 1000 g for 3 minutes. After centrifugation, supernatants were discarded, and the resulting pellets were resuspended in 1 ml PBS and transferred to 1.5 ml tubes. Each sample was washed six times with 1 ml PBS using the same centrifugation conditions as their respective protocols. Cleaned Symbiodiniaceae fractions (F2) were then resuspended in 500 µl PBS. Aliquots (200 µl) were mixed with 200 µl ATL buffer and stored at −20 °C for DNA extraction; an additional 200 µl were mixed with 200 µl RTL buffer (Qiagen, Germantown, MD, USA) and stored at −80 °C for RNA extraction. Furthermore, 100 µl of the freshly isolated Symbiodiniaceae fraction was used for bacterial isolation. An overview of the protocol is provided in Figure S1.

### Bacterial isolation and identification

Cleaned Symbiodiniaceae fractions from fresh coral samples were diluted 1:10 and 1:100 and plated on various agar media. These included marine broth 2216, 25% diluted marine broth, sterile seawater supplemented with 11 nM glucose, F/2 medium, and TCBS medium (all from MilliporeSigma, Burlington, MA, USA). Plates were incubated in the dark at 28°C for 24–72 hours, except those with F/2 medium, which were kept in a light incubator under a 12:12 h light–dark cycle at ∼50 µmol m⁻² s⁻¹. Pure bacterial isolates were maintained on marine agar at 28°C for 24 hours and then stored at room temperature.

Bacterial identification was performed by colony PCR targeting the full-length 16S rRNA gene using universal primers 27F (5′-AGRGTTYGATYMTGGCTCAG-3′) and 1492R (5′-RGYTACCTTGTTACGACTT-3′). PCR reactions contained 20 µl of the Qiagen Multiplex PCR Kit (Qiagen, Germantown, MD, USA) and 2 µl of boiled bacterial lysate (prepared in 100 µl sterile molecular-grade water). Thermocycling conditions were: 95°C for 15 min; 30 cycles of 95°C for 30 s, 57°C for 30 s, and 72°C for 60 s; followed by a final extension at 72°C for 10 min. PCR products (∼1500 bp) were visualized on 1.5% agarose gels and purified with ExoProStar (Cytiva, Marlborough, MA, USA) at 37°C for 15 min, followed by 80°C for 15 min. DNA concentrations were measured using the Qubit 4.0 Fluorometer with the 1× dsDNA HS Assay Kit (Thermo Fisher Scientific, Waltham, MA, USA). Final amplicons were stored at 4°C and submitted for Sanger sequencing to Eurofins Genomics (Louisville, KY, USA). Resulting quality-trimmed FASTA files were analyzed using the EZBioCloud 16S-based ID pipeline (Yoon et al. 2017).

### DNA and RNA isolation, amplicon library preparation and sequencing

Bacterial DNA was extracted using the DNeasy Blood and Tissue Kit (Qiagen, Germantown, MD, USA) following the manufacturer’s protocol for cultured cells, and stored at −20°C. Bacterial RNA was extracted using the RNeasy Mini Kit (Qiagen, Germantown, MD, USA) according to the protocol for cultured cells or frozen tissues, and stored at −80°C. Negative controls containing only buffers and water were included to monitor potential contamination from reagents or the laboratory environment. DNA and RNA concentrations were measured using the Qubit 4.0 Fluorometer (Thermo Fisher Scientific, Waltham, MA, USA) with the 1× dsDNA High Sensitivity (HS) Assay and RNA HS Assay kits, respectively.

To prepare 16S rRNA amplicon libraries, the V5–V6 hypervariable region of the 16S rRNA gene was amplified using primers 784F [5′-<u>TCGTCGGCAGCGTCAGATGTGTATAAGAGACAG</u>AGGATTAGATACCCTGGTA-3′] and 1061R [5′-<u>GTCTCGTGGGCTCGGAGATGTGTATAAGAGACA</u>GCRRCACGAGCTGACGAC-3′] (Andersson et al. 2008) with the corresponding Illumina adaptor overhangs underlined. Triplicate PCRs were performed for each sample using approximately 50 ng of DNA in 10 μL reactions with the Qiagen Multiplex PCR Kit and a final primer concentration of 0.5 μM. Thermal cycling conditions were: 95°C for 15 min; 30 cycles of 95°C for 30 s, 56°C for 90 s, and 72°C for 30 s; followed by a final extension at 72°C for 10 min. PCR products (∼300 bp) were visualized on 1.5% agarose gels and purified using ExoProStar (Cytiva, Marlborough, MA, USA) at 37°C for 15 min followed by 80°C for 15 min. Amplicons were quantified with Qubit and submitted to Novogene (Sacramento, CA, USA) for paired-end sequencing (2 × 301 bp) on the Illumina MiSeq platform using V3 chemistry (Illumina, San Diego, CA, USA). A total of 38 samples were sequenced, and a detailed sample list and metadata are provided in Table S1.

### 16S rRNA data analysis

Amplicon sequence variants (ASVs) were inferred using DADA2 v1.21.0 (Callahan et al. 2016). Reads with ambiguous bases or with an expected error rate >2 were removed. ASVs were denoised and merged from paired-end reads and screened for chimeras. Taxonomic classification was assigned using the SILVA database (version 138) (Quast et al. 2013), and sequence read statistics are summarized in Table S2. To minimize contamination, ASVs were filtered by removing those with a relative abundance >10% in negative controls compared to biological samples, as well as sequences affiliated with mitochondria or chloroplasts. This filtering step resulted in 1,150 ASVs retained for downstream analyses (Table S3). Beta diversity analysis was assessed using Phyloseq v1.34.0 (McMurdie and Holmes 2013), based on Euclidean distances of centered-log ratio (clr)-transformed ASV counts and visualized using principal component analysis (PCA). Differences in microbial community composition between treatments and fractions were tested using permutational multivariate analysis of variance (PERMANOVA) as implemented in the *adonis* function from the Vegan package v2.5 (Oksanen et al. 2007). For alpha diversity, observed ASV richness was calculated after rarefying all samples to the lowest read count (n = 6,597) using the *rarefy* function from the GUnifrac package (Chen and Chen 2018) and the *diversity* function from Vegan. Differences in alpha diversity were tested using a linear model fitted with the *lm* function, followed by analysis of variance (ANOVA) to assess the effects of explanatory variables. Post hoc pairwise comparisons were conducted using the *emmeans* function, which estimates marginal means based on the fitted model. P-values for pairwise tests were adjusted for multiple comparisons using the false discovery rate (FDR) method.

Relative abundances of the top 20 most abundant ASVs were visualized with ggplot2 v3.3.5 (Wickham, Chang, and Wickham 2016). Differentially abundant ASVs between F2 and F1 fractions were identified using the Analysis of Composition of Microbiomes with Bias Correction (Lin and Peddada 2020). The function *ANCOM-BC* v2.4.0 was applied to ASV absolute counts to identify ASVs enriched in F2 compared to F1 from each group of samples but also in an overall comparison irrespective of treatment. An ASV was considered significantly enriched when FDR-adjusted p-values <0.05. The number of differentially abundant ASVs were displayed in bar plots and lollipop plots that include their log fold change across F1 and F2 fractions. Overlapping ASVs were displayed in radar plots using the package fmsb v0.7.6.

### qPCR validation of Symbiodiniaceae and Vibrio enrichment across coral species

Complementary DNA (cDNA) was synthesized from RNA using the High-Capacity cDNA Reverse Transcription Kit (Qiagen, Germantown, MD, USA) according to the manufacturer’s instructions and stored at −20°C until further processing. Quantitative PCR (qPCR) was used to assess the relative enrichment of Symbiodiniaceae and *Vibrio* compared to coral and bacterial cells, respectively, by targeting taxon-specific gene markers. The mitochondrial 16S gene of the coral host was amplified using primers Sponge_16S_F [5’ NGAGTACTGTRAAGGAAAGYTG 3’] and Sponge_16S_R [5’ AGATCACTTGGYTTCGGG 3’] (Alexander et al. 2020). The ITS2 gene of Symbiodiniaceae was targeted using primers Symb_F [5’ GAATTGCAGAACTCCGTGAACC 3’] (Hume et al. 2018) and Symb_R [5’ AGCACTGAAGCAGACATACTCTCAG 3’] (Meistertzheim et al. 2019). The 16S rRNA gene was amplified as a positive control of the reactions using primers 784F [5’ AGGATTAGATACCCTGGTA 3’] and 1061R [5’ CRRCACGAGCTGACGAC 3’] (Andersson et al. 2008). Members of the genus *Vibrio* were amplified using the primers 567F [5’ GGCGTAAAGCGCATGCAGGT 3’] and 680R [5’ GAAATTCTACCCCCCTCTACAG 3’] (Thompson et al. 2004).

qPCR reactions (10 μL total volume) contained 0.5 μL each of forward and reverse primers (final concentration: 0.5 μM), 5 μL AzuraQuant Green Fast qPCR Mix LoRox (Azura Genomics, Raynham, MA, USA), 2 μL sterile PCR-grade water, and 2 μL of DNA or cDNA template (∼100 ng). Reactions were performed on the LightCycler 96 system (Roche Diagnostics, Indianapolis, IN, USA) under the following cycling conditions: initial denaturation at 95°C for 3 min; 40 cycles of 95°C for 20 s, 55°C for 1 min, and 72°C for 20 s; followed by a high-resolution melt curve with steps at 95°C for 1 min, 40°C for 1 min, 65°C for 1 s, and 97°C for 1 s. Each sample was run in technical triplicates for the coral and Symbiodiniaceae primer sets, while a single reaction was used with the bacterial primers as a positive control. Amplification and melt curve data were analyzed using the LightCycler 96 software.

Algal enrichment was calculated by first determining the mean Cq (quantification cycle) values from qPCR technical replicates for both algal and coral primers in each sample. The difference in Cq values between algae and coral (ΔCt) was computed for each sample, and fold change was calculated as 2 raised to the negative ΔCt, representing the relative abundance of algae to coral. To assess enrichment due to fractionation, fold changes from centrifugation protocols (P1, P2) were normalized by the fold change in the unfractionated samples, yielding a fold enrichment value that reflects the increase in algal abundance relative to unfractionated controls. *Vibrio* enrichment was calculated in relation to overall bacterial abundance using the approach, allowing us to evaluate the relative increase of *Vibrio* compared to the total bacterial community in each fraction.

To validate our results in additional species, three clonal fragments each of the coral species *Stylophora pistillata and Pocillopora damicornis,* were purchased from the World Wide Corals aquaria facility in Florida and frozen upon arrival. Fractionation was performed using centrifugation protocol P1, and RNA extraction, cDNA synthesis, and qPCR were conducted in duplicates as described above.

To evaluate the relationship between *Vibrio* and algal enrichment, a Pearson correlation analysis was performed using ggpubr v.0.6.0 (Kassambara 2018). Prior to analysis, potential outliers were assessed using Cook’s distance derived from a linear model (lm(Algal Enrichment ∼ Vibrio Enrichment)). The threshold for influential points was set at 4/(*n*-2), where *n* is the number of observations. One influential point with the highest *Vibrio* enrichment value was identified and removed from the dataset to reduce leverage effects on the correlation. The scatter plot was generated using the *ggscatter* function, including a fitted regression line and 95% confidence interval. Pearson’s correlation coefficient (R) and associated p-value were displayed on the plot.

### Genome sequencing

Over 20 bacteria strains identified as *Vibrio* were cultured in 10 ml of sterile marine broth 2216 NutriSelect Plus, (Sigma-Aldrich, St. Louis, MO, USA) for 24 h. The cultures were then centrifuged at 3,234 g for 15 min, the supernatant was discarded, and resulting pellets were frozen at −20°C until DNA extraction. Genomic DNA was extracted using the DNeasy Blood & Tissue Kit (Qiagen, Germantown, MD, USA), following the manufacturer’s protocol with a minor modification: 8 µL of RNase A (Qiagen, Germantown, MD, USA) were added to each sample prior to the addition of Proteinase K and Buffer AL. Samples were incubated at 56 °C and 700 rpm for 1 h. DNA concentrations were measured using both the Qubit Fluorometer 4.0 (Thermo Fisher Scientific, Waltham, MA, USA) and the NanoDrop Lite Plus (Thermo Fisher Scientific). Samples with 260/280 or 260/230 ratios below 1.8 were rewashed starting from the ethanol wash step in the manufacturer’s protocol.

DNA libraries were prepared using the Rapid Barcoding Kit 96 V14 (Oxford Nanopore Technologies, Oxford, UK) and sequenced on a GridION platform using MinION flow cells and Flongles (R10.4.1 chemistry) following the manufacturer’s instructions and community guidelines.

### Genomics analysis of Vibrio strains

Raw sequencing data for each barcode were consolidated into a single multi-FASTA file. De novo genome assemblies were performed using Flye v2.8.1 (Kolmogorov et al. 2019), and assembly quality metrics were assessed with QUAST v5.0.2 (Gurevich et al. 2013). Genome completeness and contamination were evaluated using the *lineage_wf* pipeline from CheckM v1.2.2 (Parks et al. 2015). Initial taxonomic classification of isolates was performed using the *classify_wf* module in GTDB-Tk v2.1.1 (Chaumeil et al. 2019), which assigns taxonomy based on Average Nucleotide Identity (ANI) relative to GTDB reference genomes. Closest matching reference genomes were retrieved from NCBI via the GTDB advanced search. Phylogenomic relationships among the 11 *Vibrio* genomes and eight closely related type strains, including a member of *Photobacterium* as an outgroup, were inferred using the *de_novo_wf* pipeline of GTDB-Tk. The tree was based on a concatenated alignment of 119 bacterial marker genes, comprising 5,036 amino acid positions. The resulting maximum likelihood phylogeny was visualized using <u>Phylo.io</u>.

Gene prediction and functional annotation were performed using Prokka v1.13 (Seemann 2014), which provides genome-wide annotation including identification of coding sequences, tRNAs, and rRNAs. A literature review was conducted to identify genes associated with virulence in marine *Vibrio* species, as well as genes broadly implicated in beneficial host-microbe interactions. The presence or absence of these genes was mapped onto a phylogenetic tree using ggtree v3.10.1 (Yu et al. 2017), based on the phylogenetic relationships inferred through GTDB-Tk. The genomes are available through Zenodo (Cardenas 2025).

### Comparison of ASVs to 16S rRNA Sequences from Vibrio Genomes

Sequences of all ASVs classified as *Vibrio* (n = 31) and the 16S rRNA gene sequences from 11 *Vibrio* isolate genomes were analyzed. Full-length 16S rRNA gene sequences from *Vibrio* genomes were digitally PCR-trimmed in silico using Cutadapt v5.1 (Martin 2011) to match the ASV amplicon regions obtained by metabarcoding. The degenerate forward primer AGGATTAGATACCCTGGTA and reverse primer CRRCACGAGCTGACGAC (Andersson et al. 2008) were used for trimming, applying default parameters with a minimum overlap of 10 nucleotides and a maximum error rate of 0.1. Trimmed 16S sequences served as a reference database for sequence similarity searches against ASV sequences. BLAST+ v2.16.0 was used with *blastn* optimized for highly similar sequences, configured to return only the top hit per query with a minimum percent identity of 97%. Hits with ≥ 99% identity were considered indicative of close taxonomic relatedness between ASVs and reference isolates.

## RESULTS

In this study, we evaluated how different cell fractionation protocols affect the characterization of microbial communities associated with Symbiodiniaceae in clonal fragments of the coral *Acropora nobilis.* We compared centrifugation methods (P1 vs P2), nucleic acid templates (DNA vs RNA), and sample processing conditions (fresh vs frozen) to assess their impact on the bacterial profiles of Symbiodiniaceae-enriched fractions (F2) relative to unfractionated samples (F1).

### Different protocols vary in separating coral and algal cells and result in distinct microbiomes

All protocols showed increased algal-to-coral cell ratios in Symbiodiniaceae-enriched fractions (F2) compared to unfractionated samples (F1). Algal enrichment was consistently higher in RNA-based samples, with average fold increases ranging from 13.52 to 71.66 (Figure S2). In DNA-based samples, enrichment was generally below 2 when using frozen samples, while fresh samples showed highly variable enrichment, ranging from 19.51 to over 15 million-fold (Figure S2)

Unfractionated samples from all protocols resulted in microbiomes with significantly distinct composition (Table S4, Figure 1A). The largest contributor to microbiome composition were the nucleic acid type (e.g. DNA vs RNA templates) with 27.29% of the variance explained (PERMANOVA, R^2^ =0.272, p-val = 0.001), followed by the type of sample (i.e, fresh vs frozen) with 22.68% of the variance explained (PERMANOVA, R^2^ =0.226, p-val = 0.001). Most importantly, microbiomes obtained from F1 and F2 fractions were significantly different in all protocols (overall PERMANOVA comparison, R^2^ =0.1004, p-val = 0.001. Table S4 and Figure 1A-B), with largest R2 values obtained in F2 vs F1 comparisons from RNA samples (Frozen-RNA: R^2^ =0.417, p-val = 0.014. Fresh-RNA R2 = 0.401, p-val = 0.017. Table S4). Conversely, the effect of centrifugation protocol was not significant in any group sample (Table S4). Similarly, unfractionated samples from all protocols resulted in microbiomes with significantly different alpha diversity. The largest contributor to microbiome alpha diversity was nucleic acid type (ANOVA, F = 15.398, p = 0.001), followed by sample type (ANOVA, F = 5.214, p = 0.033), while distinct protocols did not affect bacterial richness (ANOVA, F = 0.278, p = 0.604. Table S5).

**Figure 1.**
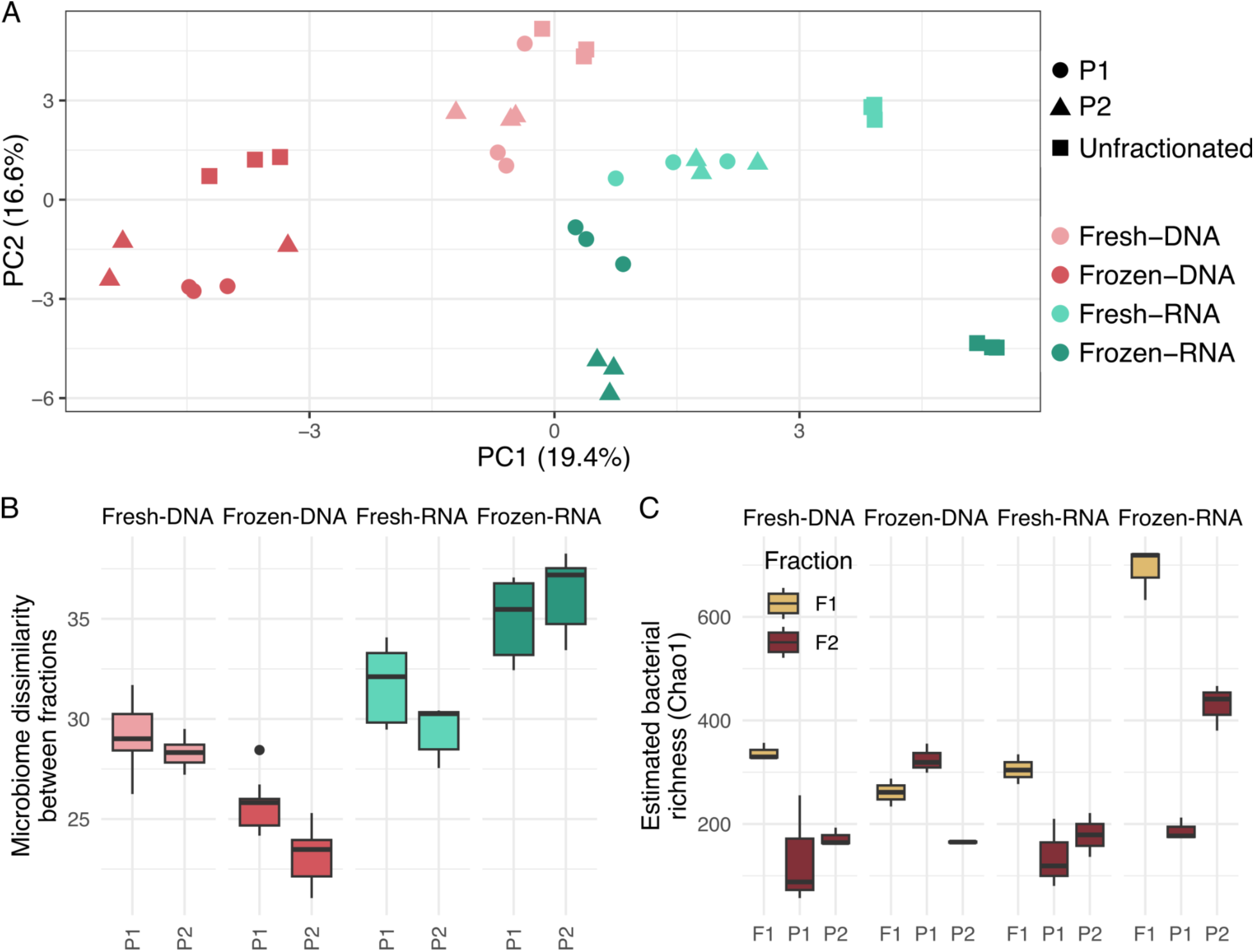
Microbiome diversity across unfractionated coral slurries and Symbiodiniaceae-enriched fractions. (A) PCA ordination plots based on Aitchison distances, showing microbial community structure across treatments (colored) and cellular fractions (shapes). (B) Microbiome dissimilarity between unfractionated (F1) and Symbiodiniaceae-enriched (F2) fractions across centrifugation protocols (P1 and P2) and sample groups. (C) Bacterial richness comparisons between F1 (yellow) and F2 (red) across protocols and sample groups.

Cell fractionation resulted in microbiomes with significantly different bacterial richness across all treatments (ANOVA, F = 6.372, p = 0.015) (Figure 1C, and Table S5). With the exception of fresh samples based on RNA template and using the centrifugation protocol P1, all Symbiodiniaceae samples had a reduction in bacterial richness (Figure 1C).

### Vibrios were significantly more abundant in Symbiodiniaceae-enriched fractions

The microbial composition of unfractionated samples (F1) revealed a dominance of *Pseudomonas* in DNA-based profiles, whereas *Endozoicomonas* predominated in RNA-based profiles (Figure 2). A notable observation was the consistently higher relative abundance of *Vibrio* in all F2 fractions compared to F1. Among the dominant taxa in F2, it is evident that sample type, nucleic acid template, and fractionation protocol influenced the enrichment of specific taxa. For example, ASV0009 (*Vibrio*) was dominant in F2 fractions from frozen samples processed using centrifugation protocol P2, whereas ASV0001 (*Vibrio*) and ASV0010 (*Vibrio*) were more abundant in frozen samples processed with P1. Conversely, ASV0004 (*Vibrio*) and ASV0005 (*Thaumasiovibrio*) were prevalent in fresh samples across protocols (Figure 2). Although ASV-specific patterns were evident, members of the genus *Vibrio* were consistently more abundant in Symbiodiniaceae-enriched fractions (Figure S3).

**Figure 2.**
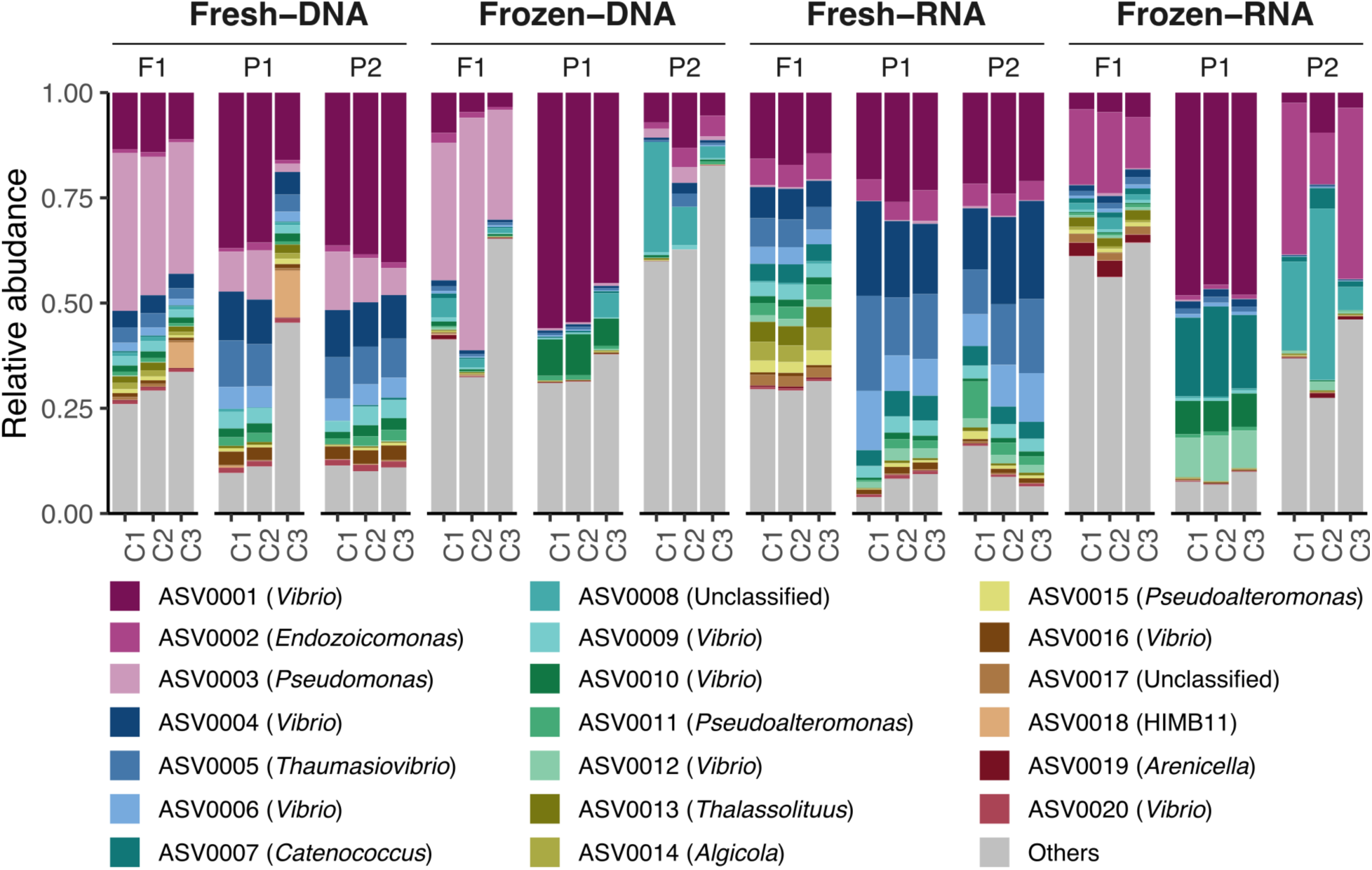
Relative abundance of top bacterial ASVs across samples. Relative abundance of bacterial 16S rRNA ASV of unfractionated (F1) and Symbiodiniacea-enriched fractions (F2), across centrifugation treatments (P1 and P2), fresh and frozen samples, and DNA and RNA templates. All other ASvs were clustered into the Others category.

Differential abundance analysis comparing F1 and F2 fractions across all treatments identified 623 significantly differentially abundant ASVs. Of these, 24 were enriched in unfractionated (F1) samples and 599 in Symbiodiniaceae-enriched (F2) fractions (Table S6). The most enriched ASVs in F2 were affiliated with the families Rhodobacteraceae (n = 88), Flavobacteriaceae (n = 74), and Vibrionaceae (n = 28) (Figure 3A, Table S6). Additional families such as Hyphomonadaceae, Rhizobiaceae, and Nitrincolaceae also exhibited both high ASV counts (n = 19, 19, and 11, respectively) and large median log fold changes in F2 fractions. Notable genera among these families include *Tenacibaculum*, *Aquimarina*, *Winogradskyella*, and *Flavobacterium* (Flavobacteriaceae); *Hellea*, *Algimonas*, and *Litorimonas* (Hyphomonadaceae); *Marinobacterium* and *Motiliproteus* (Nitrincolaceae); *Pseudahrensia* and *Cohaesibacter* (Rhizobiaceae); *Yoonia*-*Loktanella*, *Roseobacter*, *Ruegeria*, and *Marivita* (Rhodobacteraceae); and *Vibrio* and *Catenococcus* (Vibrionaceae) (Figure 3A).

**Figure 3.**
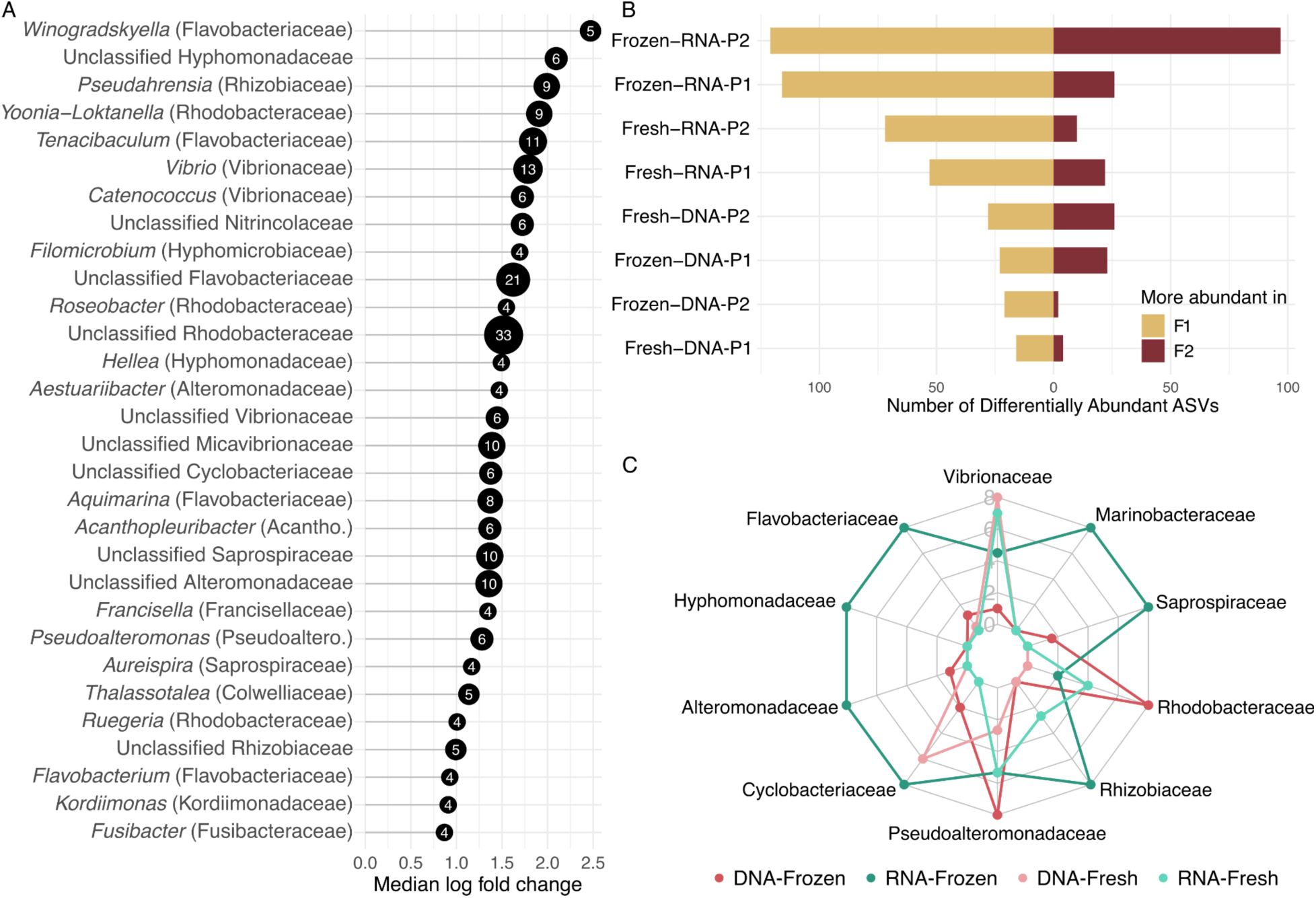
Taxonomic distribution of differentially abundant ASVs. (A) Taxonomic breakdown of ASVs significantly more abundant in the F2 fractions, based on comparisons across all fractions regardless of treatment. The x-axis displays the median log fold change for each genus, while bubble size and accompanying labels represent the number of significantly enriched ASVs within each genus. (B) Total count of differentially abundant ASVs across all treatment groups. (C) Taxonomic distribution of these differentially abundant ASVs, split by treatment. Families Pseudoalteromonadaceae and Acanthopleuribacteraceae were abbreviated as Pseudoaltero., and Acantho., respectively.

When comparing fractions within each treatment, the number of differentially abundant ASVs varied across conditions (Figure 3B, Table S7). The highest numbers were observed in frozen RNA samples (P1: n = 218; P2: n = 142), followed by fresh RNA samples (P1: n = 75; P2: n = 82). The taxonomic profiles of differentially abundant ASVs revealed a substantial proportion of shared taxa across treatments (Figures 3C, S4-5). ASV001 and ASV004 (*Vibrio*) were the most frequently enriched in Symbiodiniaceae-associated (F2) fractions across treatments, followed by other ASVs affiliated with the genera *Vibrio* and *Pseudoalteromonas* (Figure S4). The families Vibrionaceae, Flavobacteraceae, and Rhodobacteraceae were among the most consistently enriched across Symbiodiniaceae-associated fractions (Figure S5). However, when considering both enrichment and the number of shared ASVs across treatments, Vibrionaceae, Flavobacteraceae, and Hyphomonadaceae were the most prominent families (Figure 3C).

We quantified *Vibrio* enrichment in Symbiodiniaceae cell fractions from *Acropora nobilis* (Anov), *Stylophora pistillata* (Spis), and *Pocillopora damicornis* (Pdam) using qPCR. Symbiodiniaceae enrichment efficiency varied by coral species, ranging from 1–3-fold in Pdam, 1–105-fold in Spis, and 48–72-fold in Anov (Figure 4A). *Vibrio* enrichment followed a similar trend, with lower enrichment in Pdam (0.1–0.8-fold), moderate levels in Spis (2–5.3-fold), and the highest in Anov (2–7.3-fold) (Figure 4B). Overall, algal and *Vibrio* enrichment levels were strongly positively correlated (R = 0.80, p = 0.018), suggesting a consistent association between Symbiodiniaceae and *Vibrio* across coral species (Figure 4C).

**Figure 4:**
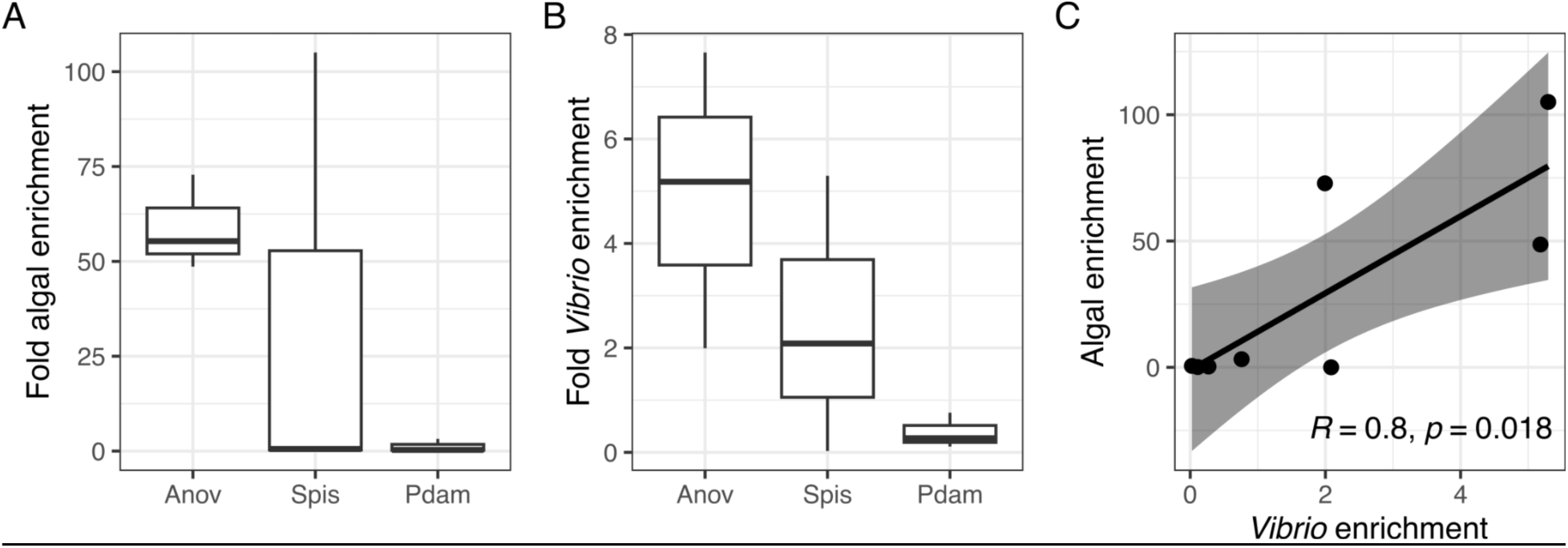
Coral species-specific enrichment of algae and *Vibro* after cell fractionation. (A) Algal and (B) *Vibrio* fold enrichment across coral species *Acropora nobilis* (Anov), *Stylophora pistillata* (Spis), and *Pocillopora damicornis* (Pdam). (C) Linear relationship between *Vibrio* enrichment and algal enrichment across samples. Pearson’s correlation coefficient (R) and associated p-value are displayed on the plot.

### Taxonomic and functional characterization of Vibrio populations enriched in Symbiodiniaceae fractions

We isolated and sequenced over 20 bacterial strains initially identified as *Vibrio*, but taxonomic classification revealed only 11 unique, non-clonal strains. These *Vibrio* genomes were nearly complete (94–100% completeness) and showed minimal contamination (0.3–2%), based on the presence of expected lineage-specific single-copy marker genes for the genus *Vibrio* (Table S8). Their GC content ranged from 44.3% to 45.8%, genome lengths from 4.8 to 6.3 Mb (Figure 5A), and predicted gene counts from 4,334 to 5,765. Ribosomal gene copy numbers varied widely, from 4 to 14 (Table S8).

**Figure 5.**
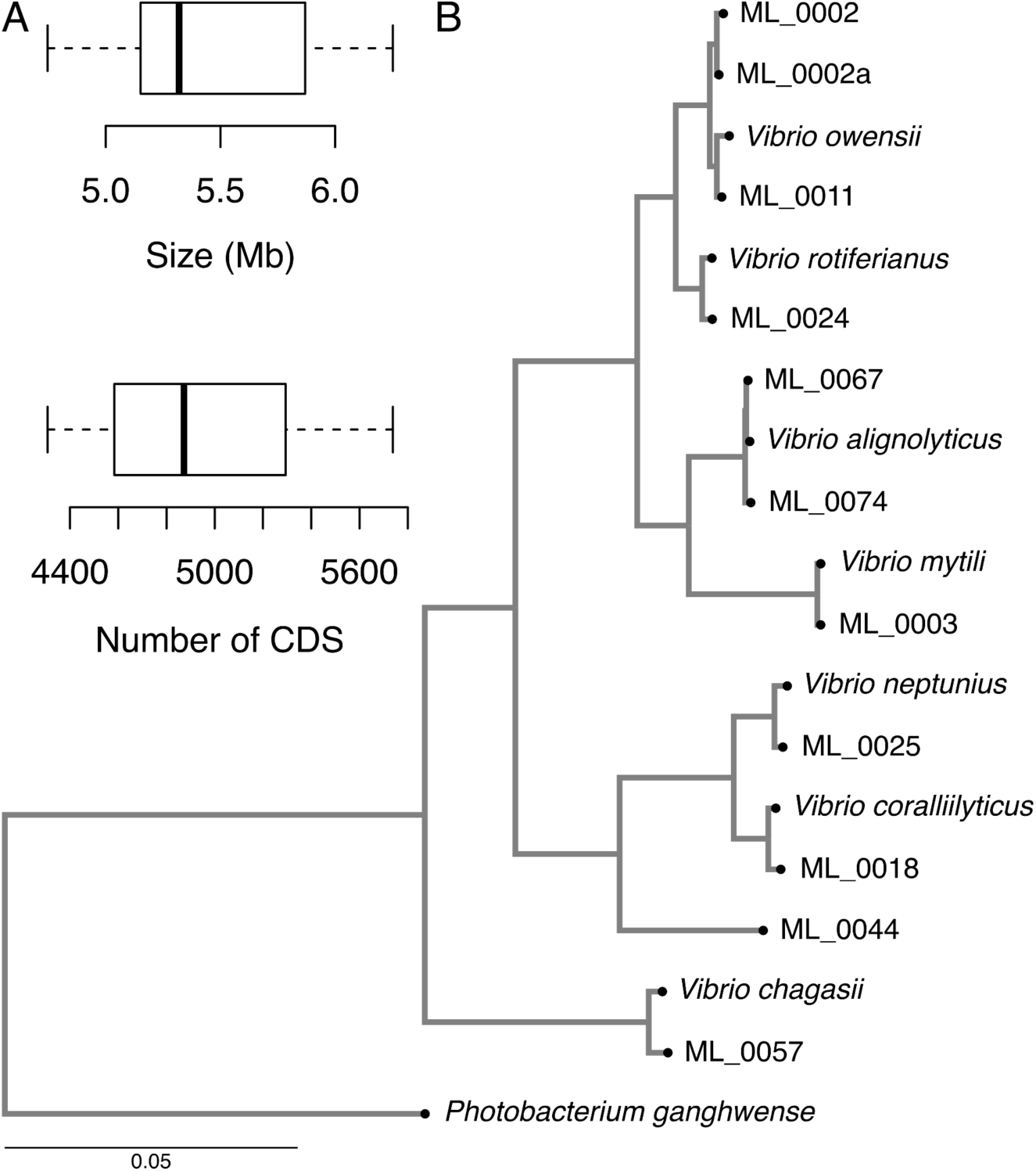
Genomic features and phylogenomic context of *Vibrio* strains isolated from Symbiodiniaceae-enriched fractions. (A) Boxplots indicating genome size and number of coding sequences (CDS). (B) Phylogenomic tree of *Vibrio* isolated from Symbiodiniaceae-enriched fractions. The phylogenomic analysis was done using GTDB-Tk using 119 molecular markers. Taxonomic classification via the classify_wf pipeline identified the closest GTDB reference species for each isolate based on Average Nucleotide Identity (ANI). A de novo phylogenetic tree was built with the de_novo_wf pipeline, including 11 *Vibrio* genomes and 7 related type strains with *Photobacterium* as an outgroup (Table S9). The analysis used 119 markers, producing a 5,036 amino acid alignment, and the tree was visualized with Phylo.io.

Average Nucleotide Identity (ANI) analysis placed the 11 Vibrio strains within three of the 51 clades recently described for Vibrionaceae (Jiang et al. 2021) (Figure 5B). Seven strains belonged to the Harveyi clade: ML_0002, ML_0002a, and ML_0011 were classified as different populations of *V. owensii*; ML_0024 as *V. rotiferianus*; ML_0067 and ML_0074 as *V. alginolyticus*; and ML_0003 as *V. mytili*. Three strains fell within the Coralliilyticus clade: ML_0025 was identified as *V. neptunius*, ML_0018 as *V. coralliilyticus*, and ML_0044 showed no close species match above the 95% ANI threshold (Table S8). The final isolate, ML_0057, belonged to the Splendidus clade and was classified as *V. chagasii*.

Analysis of the *Vibrio* genomes revealed that all strains encode multiple virulence factors (Figure 6). These include thermolabile hemolysins (present in all strains except ML_0003), the hemolysin transporter *shlB* (in ML_0018 and ML_0025), and the regulatory gene *hlyU* (present in all genomes) (Figures 6, S6; Table S10). The cytolysin gene *vvhA* was detected in ML_0018 and ML_0025, while *rtxA*, encoding RTX toxins, was found in four strains (ML_0002a, ML_0011, ML_0025, and ML_0044). Notably, the RTX-I toxin translocation system gene *apxIB* was present in all genomes. Additional virulence genes included *ltxA/B*, encoding a pore-forming toxin and its transporter (found in all strains except ML_0018 and ML_0044), and the ribosome-associated toxin gene *ratA* (present in all genomes except ML_0024).

**Figure 6.**
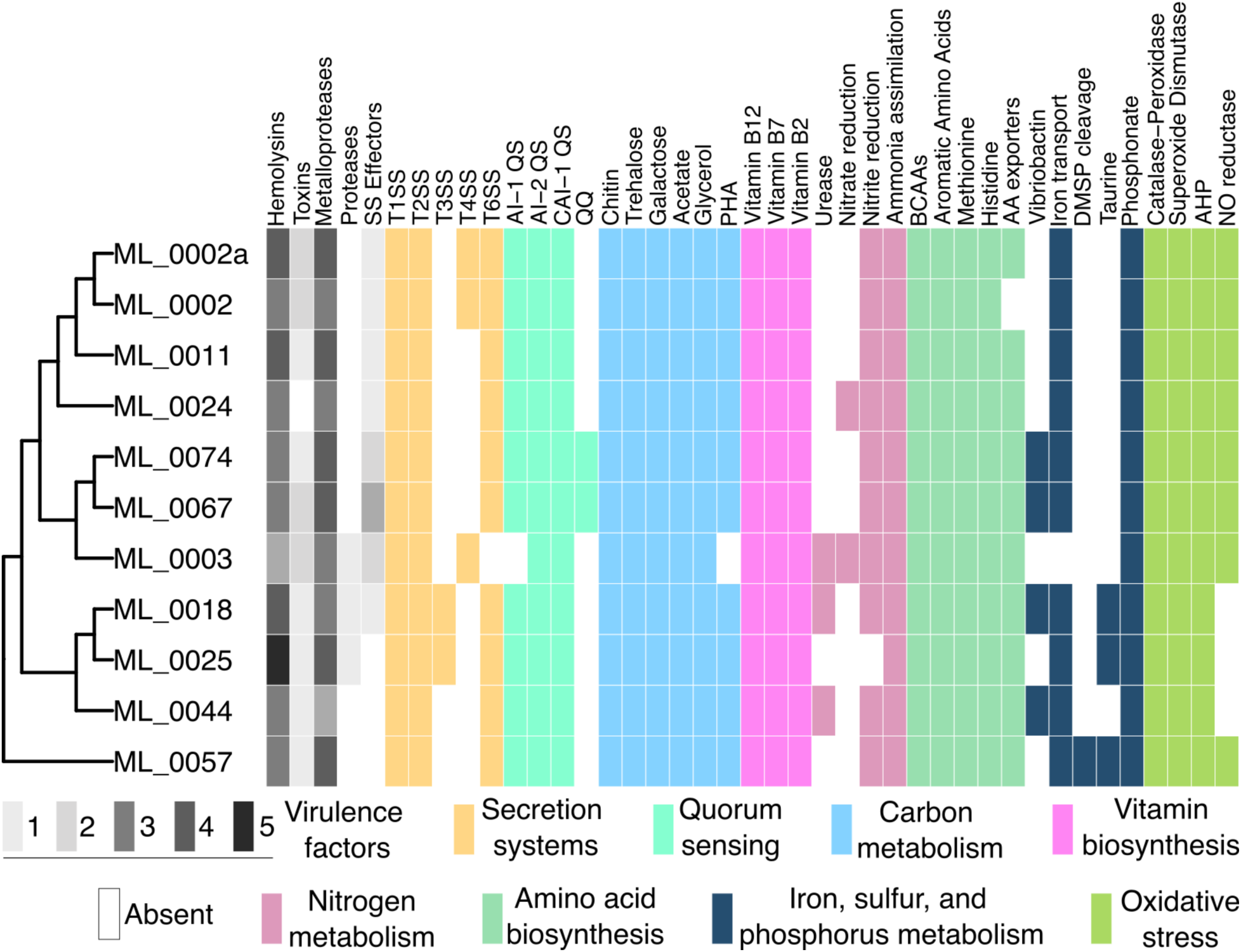
Functional traits of Symbiodiniaceae-associated *Vibrio* strains. Presence or absence of key virulence genes and genes from pathways potentially involved in beneficial symbioses. For virulence factors, the scale represents the number of distinct genes identified in each strain, while for all other categories, the plot indicates simple presence or absence. Phylogenetic relationships were inferred using the phylogenomic approach described in Figure 4, excluding reference strains.

Metalloprotease genes showed variable distribution across genomes (Figures 6, S6; Table S10). The serralysin zinc metalloprotease gene *aprA* was detected in ML_0002a, ML_0011, and ML_0057, while serralysin-like genes *prtB/C* were found in ML_0003, ML_0018, ML_0025, ML_0044, ML_0067, and ML_0074. The zinc metalloprotease gene *prtV* was present in all strains except ML_0003 and ML_0044, and *stcE* was found in ML_0002, ML_0002a, ML_0011, and ML_0024. Universally present were *tldD* and *pmbA*, which encode proteases, while *empA* (an extracellular metalloprotease) was found only in ML_0025. The gene *impA* was present in ML_0003, ML_0057, ML_0067, and ML_0074. Finally, the serine protease gene *tlp* was identified in ML_0018 and ML_0025, while *htrA* was unique to ML_0003.

Genes encoding components of the Type I, II, and VI secretion systems (T1SS, T2SS, and T6SS) were consistently present across all genomes (Figure 6; Table S10). In contrast, genes associated with Type III (spiA) and Type IV (virB) secretion systems were detected only in a subset of strains: *spiA* and *virB* were present in ML_0018 and ML_0025, and in ML_0002, ML_0002a, and ML_0003, respectively. Effector proteins linked to these secretion systems (*vopS, exsAC, hcpA,* and *tdeA*) were distributed heterogeneously among the genomes. Notably, the strains with the highest number of putative virulence factors, including multiple hemolysins, metalloproteases, toxins, and extracellular proteases, were ML_0025 (*V. neptunius*) and ML_0067 (*V. alginolyticus*), each with 14 identified factors, followed by ML_0018 (*V. coralliilyticus)*, a known Symbiodiniaceae pathogen, with 12 virulence-related genes.

Quorum sensing-related genes were widely distributed across the *Vibrio* genomes analyzed (Figure 6; Table S10). Nearly all genomes encoded key components of quorum sensing pathways, including *luxN, vanM, luxPQ, rhtB, cqsAS*, and *luxO*, indicating the presence of both Autoinducer-1 (AHL-based) and Autoinducer-2 (AI-2) communication systems. All genomes also harbored *cqsAS*, which is involved in intra-genus signaling via the (S)-3-hydroxytridecanone (CAI-1) autoinducer. Notably, the genomes of ML_0067 and ML_0074 carried the *Y2-aiiA* gene, which encodes an N-acyl homoserine lactonase capable of degrading AHL signals, potentially disrupting quorum sensing-mediated communication.

All *Vibrio* genomes analyzed shared a conserved set of genes involved in carbon acquisition, metabolism, and storage (Figures 6, S6; Table S10). Key genes for chitin degradation and utilization, including *chiAB* (chitinases), *cbpD* (chitin-binding proteins), and *chiP* (chitoporin), were universally present, along with sugar metabolism genes such as *treA/B* for trehalose import and degradation via the PTS system, *glpF* and *glpK* for glycerol uptake and metabolism, and *galK* for galactose utilization. Genes involved in acetate metabolism (*acs*) and polyhydroxyalkanoate (PHA) biosynthesis (*phaABC*) were also detected in all genomes, indicating potential for carbon storage under nutrient-limited conditions. Notably, none of the genomes carried *glcP*, a glucose transporter of the Major Facilitator Superfamily (MFS), suggesting limited capacity for direct glucose uptake (Figures 6, S6; Table S10).

Analysis of vitamin biosynthesis and transport genes revealed widespread metabolic versatility (Figures 6, S6; Table S10). Most genomes carried biosynthetic genes for vitamin B₁₂ (*cob/cbi*), particularly key late-pathway genes (*cobS, cobT, cobP, cobB*), with the remaining genomes lacking only these components. All genomes possessed vitamin B₁₂ transport genes (*btuBCDF*), indicating the ability to acquire cobalamin from external sources. Similarly, ten genomes contained the complete *bioABDF* operon for vitamin B₇ (biotin) biosynthesis, while one had only *bioA*. Vitamin B₂ (riboflavin) biosynthesis genes (*folABCD*) were also broadly conserved, with most genomes carrying *folBCD* or the full operon (Figures 6, S6; Table S10).

In terms of nitrogen metabolism, the genomes harbored an incomplete and likely non-functional set of nitrogen fixation genes (e.g., *vnfA* and partial *nif* operons), possibly acquired via horizontal gene transfer. However, all genomes encoded *napAB* and *nirD*/*nasD*, consistent with a functional assimilatory nitrate reduction pathway converting nitrate to ammonium. ML_0003 and ML_0024 additionally carried the *narGHIJ* operon, suggesting potential for respiratory nitrate reduction under low-oxygen conditions. Most genomes (except ML_0003 and ML_0025) contained *nrfABG*, key components of the DNRA pathway for nitrite reduction to ammonium, which may contribute bioavailable nitrogen to the surrounding environment. The universal presence of *glnA* supports the ability of these strains to assimilate ammonium into organic nitrogen compounds (Figures 6, S6; Table S10).

Regarding amino acid biosynthesis, all genomes encoded complete operons for branched-chain amino acids (BCAAs: isoleucine, leucine, valine via *ilvGMEDA*), aromatic amino acids (tryptophan, phenylalanine, tyrosine via *trpABCE*), and sulfur-containing amino acids (methionine via *metABCEFHE*). Most also contained the full histidine biosynthesis pathway (*hisGDHC*), with one genome missing *hisD*, potentially due to assembly or annotation issues. The branched-chain amino acid transporter gene *brnQ* was present in all genomes, while elements of the *livFGHMK* operon involved in amino acid transport were found only in a few strains, suggesting limited redundancy in amino acid export mechanisms (Figures 6, S6; Table S10).

Genes involved in iron acquisition, including siderophore production, transport, and regulation, were broadly distributed across the *Vibrio* genomes (Figures 6, S6; Table S10). Four strains (ML_0018, ML_0044, ML_0067, and ML_0074) contained the *iucABC* gene cluster, essential for the biosynthesis of the siderophore vibriobactin. Nearly all genomes carried genes associated with siderophore transport (*yusV/yfhAL*), and all strains possessed the global iron regulator *fur*, which governs siderophore production and iron uptake in response to iron availability. In contrast, genes involved in the symbiotic exchange of sulfur compounds were relatively sparse. The *dddP* gene, which encodes a DMSP lyase that breaks down dimethylsulfoniopropionate (DMSP), was found only in ML_0057, while *tauD*, associated with taurine utilization, was detected in ML_0018, ML_0025, and ML_0057 (Figures 6, S6; Table S10).

Lastly, all 11 Vibrio genomes analyzed contained key antioxidant defense genes, including *katG* (catalase-peroxidase), *ahpC* (alkyl hydroperoxide reductase subunit C), and at least one form of superoxide dismutase (*sodA* and/or *sodB*), indicating a conserved capacity to neutralize reactive oxygen species (ROS). Additionally, *norVW*, which encodes nitric oxide reductase for the detoxification of nitric oxide (NO), was present in 9 out of 11 genomes, suggesting some variability in the ability of different strains to manage nitrosative stress (Figures 6, S6; Table S10).

### Comparison between genomic and ASV 16S rRNA sequences of Vibrios

We extracted the 16S rRNA gene sequences from the *Vibrio* genomes and compared to the 31 ASVs classified as *Vibrio* (Table S11). Of these 31 ASVs, 10 matched most closely to ML_0057 (*V. chagasii*), 9 to ML_0018 (*V. coralliilyticus*), 9 to ML_0003 (*V. mytili*), 2 to ML_0044 (*Vibrio* sp.), and 1 to ML_0025 (*V. neptunius*). Similarly, among the 16 Vibrio ASVs that were significantly enriched in one or more Symbiodiniaceae-enriched fractions, 6 matched ML_0003 (including ASV0001 with 100% sequence similarity), 3 (ASV0004, ASV0006, and ASV0116) matched ML_0018, 3 matched ML_0057 (ASV0016, ASV0020, and ASV0025), and 1 matched ML_0044 (ASV0038).

## DISCUSSION

The microbiome associated with in hospite Symbiodiniaceae (within the coral host) may play a critical but understudied role in coral health and resilience. While bulk sequencing of coral holobionts has revealed diverse bacterial communities, it lacks the spatial resolution to identify microbes directly interacting with Symbiodiniaceae inside the host. Identifying these closely associated bacteria is key to understanding their potential influence on coral physiology, nutrient exchange, and stress tolerance (Matthews et al. 2020; Garrido et al. 2021). However, dissecting host microcompartments remains technically challenging, and minimizing contamination from surrounding coral tissue is essential for accurately characterizing Symbiodiniaceae-associated microbiomes. In this study, we assessed how different steps in the sampling processing influence the composition of microbial communities associated with Symbiodiniaceae-enriched fractions as a proxy for identifying prevalent members of Symbiodiniaceae microbiomes. Using clonal fragments of *Acropora nobilis*, we compared centrifugation methods (P1 vs. P2), nucleic acid templates (DNA vs. RNA), and sample processing conditions (fresh vs. frozen) to evaluate their impact on the bacterial profiles of Symbiodiniaceae-enriched fractions (F2) relative to unfractionated samples (F1).

### Sample processing has large effects on coral microbiome composition

Coral microbiome composition has been extensively studied, and sample processing and library preparation methods are known to significantly influence observed bacterial diversity (Silva, Epstein, and Vega Thurber 2022; Pratte and Kellogg 2021; Bergman et al. 2022). Among the variables tested, the most pronounced effect was observed between DNA- and RNA-based library preparations. RNA-based community profiling has emerged as a useful approach to target active microbial members, distinguishing them from DNA-based profiles that include both active and inactive or dead cells (Li et al. 2017). While not a direct measure of viability, RNA provides a proxy for identifying generally viable or metabolically active community members (Ya Wang et al. 2023). Our results show that DNA-based unfractionated coral microbiomes are dominated by *Pseudomonas*, while RNA-based profiles are dominated by *Endozoicomonas*. This suggests *Endozoicomonas* is an active member of the coral-associated microbiome. Although direct metatranscriptomic evidence for *Endozoicomonas* activity in coral tissues is lacking, several RNA-based studies have reported it as a prevalent member of the community (Savary et al. 2021; Voolstra et al. 2021; Ziegler et al. 2017). Additionally, others have shown that *Endozoicomonas* can mount a pronounced transcriptomic response to coral tissue extracts, supporting its potential functional role (Pogoreutz et al. 2022).

We also found that protocol choice significantly affected fractionation efficiency. On average, frozen-based protocols produced more consistent results with lower variability, likely because fresh samples yield excess mucus that hinders clean separation, an issue others have addressed by freezing (Wallace et al. 2024). However, freezing can cause some cells to lyse, reducing overall biomass yield. Despite this trade-off, combining sample freezing, slower and longer centrifugation steps (P1), and RNA-based profiling produced the most consistent results. This was evidenced not only by higher algal-to-coral cell ratios but also by greater microbiome dissimilarity and reduced richness relative to unfractionated samples. Still, these outcomes may vary across coral species, and sample characteristics should be evaluated on a case-by-case basis.

### Vibrios and other key players are consistently enriched in Symbiodiniaceae fractions

In this study, some of the most conspicuous members of the in hospite Symbiodiniaceae microbiomes belonged to the family Flavobacteriaceae (phylum Bacteroidota), including genera such as *Winogradskyella*, *Tenacibaculum*, *Aquimarina*, and *Flavobacterium*. Previous research has shown that members of the class Flavobacteriia maintain a close spatial association with Symbiodiniaceae and may even occur intracellularly within the algal endosymbiont (Maire et al. 2021; Hill et al. 2024). Members of Flavobacteriaceae are also commonly found in Symbiodiniaceae cultures (Díaz-Almeyda et al. 2022), and some strains (specifically *Muricauda lutaonensis* CC-HSB-1) have been shown to enhance thermal and light tolerance of cultured Symbiodiniaceae via the production of the antioxidant carotenoid zeaxanthin (Motone et al. 2020). Within the Proteobacteria, members of the Alphaproteobacteria class have also been previously found in close spatial proximity to Symbiodiniaceae cells (Maire et al. 2021), particularly taxa such as Rhizobiales (now reclassified as Hyphomicrobiales), *Bradyrhizobium*, Rhodobacteraceae, and Sphingomonadaceae (Gardner, Leggat, and Ainsworth 2023; Hill et al. 2024). In this study, we corroborate the presence of Hyphomicrobiales with a dominance of members from the Rhizobiaceae, Hyphomicrobiaceae, and Stappiaceae families. These groups have been classically associated with terrestrial plant environments, but are increasingly detected in phytoplankton-rich aquatic systems (Camacho et al. 2016; Z. Wang et al. 2024). Rhizobiaceae are typically nitrogen-fixing symbionts (Tschitschko et al. 2024), while Hyphomicrobiaceae are known to metabolize methylated compounds such as methanol, which is commonly released by phytoplankton (Dixon, Beale, and Nightingale 2011). Stappiaceae, particularly *Labrenzia* spp., have been shown to form persistent associations with cultured Symbiodiniaceae and exhibit algal growth-promoting abilities (Lawson et al. 2018; Matthews, Hoch, et al. 2023). However, in our study, *Labrenzia* members were enriched in both fractionated and unfractionated samples, preventing us from drawing firm conclusions about their specific spatial association in hospite.

We also confirmed the presence of Rhodobacteraceae in Symbiodiniaceae-enriched fractions, specifically several members of the Yoonia-Loktanella group, as well as the genera *Roseobacter*, *Ruegeria*, and *Marivita*. Rhodobacteraceae are widely found in coral-associated microbiomes (Sweet et al. 2021), in association with free-living phytoplankton (Focardi et al. 2025; Shibl et al. 2020), and Symbiodiniaceae cultures (Camp et al. 2020; Maire et al. 2021). Consistent with Hill et al. (2024), we also detected members of the Sphingomonadaceae family, which have previously been proposed to be intracellular associates of Symbiodiniaceae in culture (Maire et al. 2021).

Similar to previous research (Hill et al. 2024; Maire et al. 2021), we found members of the Gammaproteobacteria significantly enriched in Symbiodiniaceae fractions. Notably, *Marinobacter*, similar to *Labrenzia*, has been identified as part of the core microbiome of cultured Symbiodiniaceae and has demonstrated algal growth-promoting capabilities (Lawson et al. 2018; Matthews, Hoch, et al. 2023). While Labrenzia is less commonly observed in bulk coral microbiomes, *Marinobacter* is frequently found in coral skeletons (Cárdenas et al. 2022), larval stages (Sharp, Distel, and Paul 2012), and is one of the few taxa shown to be vertically transmitted via gametes (Cárdenas et al. 2025). These patterns suggest that *Marinobacter* is a consistent and potentially key member of Symbiodiniaceae-associated microbiomes in hospite.

Members of the genus *Vibrio* were the most conspicuous taxa in Symbiodiniaceae-enriched fractions. Although often regarded as mucus-associated (Rubio-Portillo, Robertson, and Antón 2024), some evidence suggests that *Vibrio* spp. can closely associate with Symbiodiniaceae. For example, (Maire et al. 2021) reported the presence of Gammaproteobacteria in algal fractions, though without species-level resolution. (Hill et al. 2024), found Vibrionaceae to dominate across all SIP fractions, and consistent with our findings, *Vibrio* was the most abundant genus across Symbiodiniaceae-enriched fractions, though this was not discussed in their study.

The *Vibrio shiloi–Oculina patagonica* model of coral bleaching provides mechanistic insights into *Vibrio*–Symbiodiniaceae interactions. *V. shiloi* first adheres to β-D-galactopyranoside receptors on the coral surface (Toren et al. 1998), invades the epidermis (Banin et al. 2000), transitions to a viable but non-culturable (VBNC) state (Israely, Banin, and Rosenberg 2001), and ultimately produces toxins that impair photosynthesis and kill Symbiodiniaceae (Y. Ben-Haim et al. 1999). While many vibrios appear confined to coral mucus, this model and recent research, suggest that under certain conditions, they can colonize deeper tissues, including gastrodermal cells (Gavish et al. 2021; Tang et al. 2020; Gibbin et al. 2018). Vibrios are also known to be facultative intracellular symbionts in other systems (de Souza Santos and Orth 2014; Abd et al. 2007). However, a recent study characterizing gastrodermal cavity microbiomes did not detect vibrios (Bollati et al. 2024), indicating that such associations may be context-dependent.

We isolated diverse *Vibrio* species from Symbiodiniaceae-enriched fractions, spanning 3 clades (Harveyi, Coralliilyticus, and Splendidus), and including 7 species: *V. owensii*, *V. rotiferianus*, *V. alginolyticus*, *V. mytili, V. neptunius*, *V. coralliilyticus*, and *V. chagasii*. Comparisons between ASVs and isolate-derived 16S rRNA sequences revealed identical or near-identical matches with ASVs significantly enriched in Symbiodiniaceae fractions, confirming that several of our isolates (e.g., *V. coralliilyticus, V. mytili,* and *V. chagasii*) represent dominant populations associated with Symbiodiniaceae. However, because ASVs represent short (∼300 bp) fragments, these matches are strong indicators but not definitive evidence of genomic identity, particularly given the limited taxonomic resolution of 16S in this group.

### Symbiodiniaceae-associated Vibrios are highly virulent but also harbor putative beneficial traits

All of these species have previously been found in corals, and most have been prominently characterized as causative agents of coral diseases and bleaching (Ushijima et al. 2012; Cervino et al. 2008; Zhenyu et al. 2013; Y. Ben-Haim et al. 2003). However, the mechanisms of *Vibrio* infection in corals remain largely unknown, with the exception of *V. coralliilyticus*, whose virulence is better understood (W. Wang et al. 2022; Lydick et al. 2025; Mass et al. 2024). It is generally accepted that vibrios are commensal associates that shift toward a pathogenic lifestyle under elevated temperatures (Sheikh et al. 2022; Vezzulli et al. 2015; Vezzulli, Colwell, and Pruzzo 2013). In corals, this temperature-dependent virulence switch has been demonstrated for *V. coralliilyticus* and *V. shiloi* (W. Wang, Tang, and Wang 2024; Kimes et al. 2012; Frydenborg et al. 2014) and this pattern likely extends to other Vibrio species. In addition to temperature, factors such as cell density (Zhang et al. 2023), iron availability (Payne, Mey, and Wyckoff 2016), and host-derived cues (Johnson 2013), modulate virulence in vibrios. Nevertheless, the nature of the interactions between vibrios and their coral and algal hosts, whether commensal or beneficial, remains unclear.

The presence of multiple virulence-associated genes in these *Vibrio* populations, including cytolysins (*vvhA, rtxA, hlyA*), zinc metalloproteases (*aprA, prtV, empA*), and secretion system effectors (*vopS, hcpA*), indicates a highly virulent phenotype capable of mediating antagonistic interactions between Symbiodiniaceae and Vibrio. These gene products are known to disrupt host cell membranes (Yuan, Feng, and Wang 2020; Lee et al. 2007), degrade extracellular proteins (Osei-Adjei, Huang, and Zhang 2018; Denkin and Nelson 2004), and interfere with cellular processes through direct injection of toxic effectors (Williams et al. 1996). In particular, zinc metalloproteases have been shown to play a central role in coral bleaching pathology by disrupting photosystem II, leading to photoinhibition and reduced algal viability (Sussman et al. 2009). In early studies of the coral pathogen *V. coralliilyticus*, a zinc metalloprotease was purified, and its application to coral fragments resulted in dose-dependent tissue damage within 18 hours at 27–28°C (Yael Ben-Haim, Zicherman-Keren, and Rosenberg 2003). However, a ΔvcpA mutant of *V. coralliilyticus* strain P1 caused similar levels of mortality in two animal models and in Symbiodiniaceae, demonstrating that its pathogenicity is not dependent on a single virulence factor (Santos et al. 2011). Additionally, structural and functional components of the type VI secretion system (T6SS) are important virulence factors in *V. coralliilyticus*, yet, similarly to ΔvcpA, inactivation of T6SS2 did not completely abolish toxicity in the host (Mass et al. 2024).

The widespread presence of multiple quorum sensing (QS) systems suggests that these *Vibrio* strains are well-equipped to regulate density-dependent behaviors, including virulence, as previously demonstrated (Lydick et al. 2025; Ushijima et al. 2018). Interestingly, the detection of *Y2-aiiA* in two genomes suggests a potential for quorum-quenching activity, which could serve as a competitive strategy against neighboring vibrios or other bacteria, particularly within the spatially constrained environment of the symbiosome. While *Vibrio* species are prolific producers and users of acyl-homoserine lactones (AHLs) for QS, to our knowledge, there is no previous evidence that they naturally produce quorum-quenching enzymes such as AHL lactonases or acylases.

The presence of a diverse array of virulence factors suggests that these *Vibrio* strains are well-equipped to colonize, outcompete, and damage both microbial competitors and eukaryotic hosts. However, their high prevalence in Symbiodiniaceae-enriched fractions of healthy corals, both in this study and previous work (Hill et al. 2024), suggests that their densities must be tightly regulated by host or microbiome factors, or that they may also serve beneficial roles, including participation in nutrient cycling. Notably, members of the Vibrionaceae represent the largest fraction of phytoplankton-associated generalist bacteria, as well as a substantial proportion of specialists, highlighting their importance as microalgal associates (Focardi et al. 2025).

Here, we highlight the potential of *Vibrio* strains to support symbiosis with Symbiodiniaceae through diverse metabolic functions that facilitate nutrient exchange and may enhance coral holobiont resilience. In terms of carbon metabolism, the consistent absence of glucose transporter genes (e.g., glcP) suggests these bacteria do not rely on glucose, a major photosynthate of Symbiodiniaceae (Burriesci, Raab, and Pringle 2012). Instead, their genomes indicate adaptations for alternative host-derived carbon sources, including trehalose (*treA, treB*), galactose (*galK*), glycerol (*glpF, glpK*), and acetate (*acs*), all of which are known products of Symbiodiniaceae and coral metabolism (Suescún-Bolívar, Traverse, and Thomé 2016; Hagedorn et al. 2015; Patton, Abraham, and Benson 1977). A complete chitin degradation pathway (*chiAB, cbpD, chiP*) further supports utilization of chitin, a coral host-derived substrate previously shown to modulate gene expression in *Vibrio* (Giubergia et al. 2017).

With regard to nitrogen cycling, the presence of genes for dissimilatory nitrate reduction to ammonium (DNRA) and ammonium assimilation via glnA suggests potential roles in nitrogen provisioning and competition. This is particularly relevant given that nitrogen-limiting conditions are essential for maintaining a stable and beneficial coral-algal symbiosis (Rädecker et al. 2021). Although several Vibrio species are known diazotrophs (Chimetto et al. 2008; Urdaci, Stal, and Marchand 1988), the lack of nitrogen fixation genes in these strains suggests they do not contribute new nitrogen to the Symbiodiniaceae. In addition to carbon and nitrogen metabolism, these *Vibrio* genomes encode biosynthetic pathways for several B vitamins, including B1 (thiamine), B2 (riboflavin), B6 (pyridoxal), and B12 (cobalamin). These vitamins are essential cofactors for Symbiodiniaceae, which are known to be cobalamin auxotrophs (Croft et al. 2005; Matthews et al. 2020).

Siderophore biosynthesis genes were present only in a subset of *Vibrio* genomes, suggesting that siderophore-mediated iron scavenging is not a universal trait among these strains. It is plausible that non-producers may benefit from siderophores released by other strains. While siderophore production in *Vibrio* has been associated with virulence (Balado et al. 2018; Litwin, Rayback, and Skinner 1996), similar to plant-associated bacteria, it may also confer benefits to algal hosts by increasing iron bioavailability (Scavino and Pedraza 2013).

Regarding sulfur metabolism, only a few strains encoded ddd lyases involved in the degradation of dimethylsulfoniopropionate (DMSP), an organosulfur compound produced by Symbiodiniaceae and other phytoplankton species (Steinke et al. 2011). DMSP degradation enables bacteria to access sulfur and carbon sources (Raina et al. 2010, 2009), and has been associated with the suppression of opportunistic pathogens and promotion of beneficial traits (Raina et al. 2016; Santoro et al. 2021).

Finally, genes involved in reactive oxygen species (ROS) detoxification were consistently present across *Vibrio* genomes, suggesting a widespread capacity to manage oxidative stress. This trait is commonly found in beneficial bacteria and is important in protecting Symbiodiniaceae from oxidative damage (Dungan et al. 2021). Moreover, the high prevalence of nitric oxide (NO) reductase genes points to a potential role in modulating NO-based signaling with coral and algal hosts. This signaling mechanism has been implicated in symbiotic regulation in animals, plants, and algae (Yanling Wang and Ruby 2011; Astier et al. 2021), and is particularly relevant in the context of thermal stress, which can induce NO accumulation in Symbiodiniaceae (Bouchard and Yamasaki 2008).

## CONCLUSIONS

In this study, we assessed how different steps in the sampling process influence the composition of microbial communities associated with Symbiodiniaceae microbiomes. Despite some differences in taxonomic profiles between protocols, all approaches consistently revealed a conserved core of bacterial taxa closely associated with Symbiodiniaceae. Prominent among these were members of the Vibrionaceae, Flavobacteriaceae, and Rhodobacteraceae, as well as *Marinobacter* genus, commonly linked to mutualistic roles in other microalgal systems. Notably, *Vibrio* species were consistently enriched in Symbiodiniaceae fractions isolated from healthy corals, warranting closer examination of their functional potential. Genomic analyses of these *Vibrio* populations revealed a dual repertoire of genes, those associated with virulence, including cytolysins, zinc metalloproteases, and T6SS effectors, and others linked to beneficial functions such as nutrient metabolism, oxidative stress resistance, and siderophore production. This apparent functional plasticity suggests that *Vibrio* spp. may adopt context-dependent roles within the coral-algal symbiosis, shaped by host factors, environmental conditions, and microbial community interactions. While their capacity for pathogenicity remains a concern under stress, their widespread occurrence in healthy tissues and metabolic versatility indicate that some lineages may confer ecological advantages to their hosts. Overall, these findings expand our understanding of coral holobiont complexity and highlight the need to reframe coral-associated *Vibrio* not solely as pathogens but as dynamic symbionts with the potential to influence holobiont stability and resilience.

## DATA AVAILABILITY

Scripts used are deposited in the GitHub repository https://github.com/ajcardenasb/Symbiodiniaceae-associated_vibrios/. Raw sequencing data are deposited in the NCBI Sequence Read Archive (SRA) under BioProject PRJNA1265228 (https://www.ncbi.nlm.nih.gov/bioproject/PRJNA1265228) and BioSamples SAMN48590778 to SAMN48590815.*m* Genome assemblies are available at https://doi.org/10.5281/zenodo.16754531.

## AUTHOR CONTRIBUTIONS

KJ and AC designed the experiment. KJ, CK, and AE processed coral samples. KJ, MS, and CK purified and identified bacterial isolates. CK and AS prepared and sequenced libraries. CK and AE conducted qPCRs. AS and AC analyzed bacterial diversity and phylogenomic data. AC and LS analyzed virulence gene annotations. AC analyzed other functional categories. AC wrote the manuscript with input from all co-authors.

## ACKNOWLEDGMENTS

We thank Dr. John Bracht for granting access to the qPCR thermocycler and Oxford Nanopore sequencing platform

---

**Figure S1.**
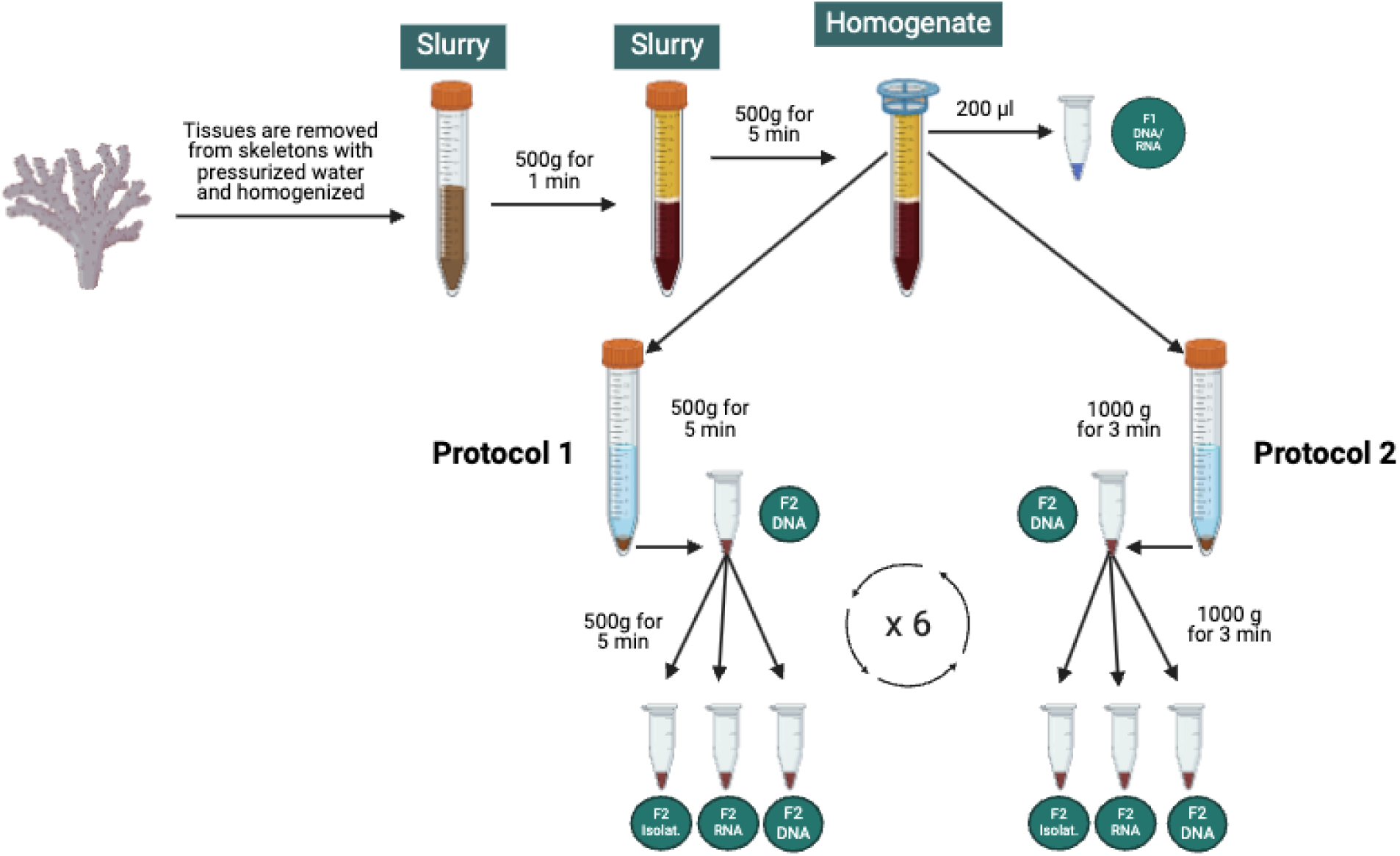
Experimental design for generating Symbiodiniaceae-enriched fractions from fresh and frozen coral fragments. Coral tissues were removed using pressurized air in phosphate-buffered saline (PBS). Samples were either kept unfractionated (F1) or processed to obtain Symbiodiniaceae-enriched fractions (F2) using one of two centrifugation protocols: P1 (500 × g for 5 min) or P2 (1000 × g for 3 min). Each fraction was washed six times and resuspended in 1 mL of PBS. F2 samples derived from both fresh and frozen coral tissue were used for RNA and DNA extractions, while fresh F2 material was also used for bacterial isolation. Image created using BioRender.

**Figure S2.**
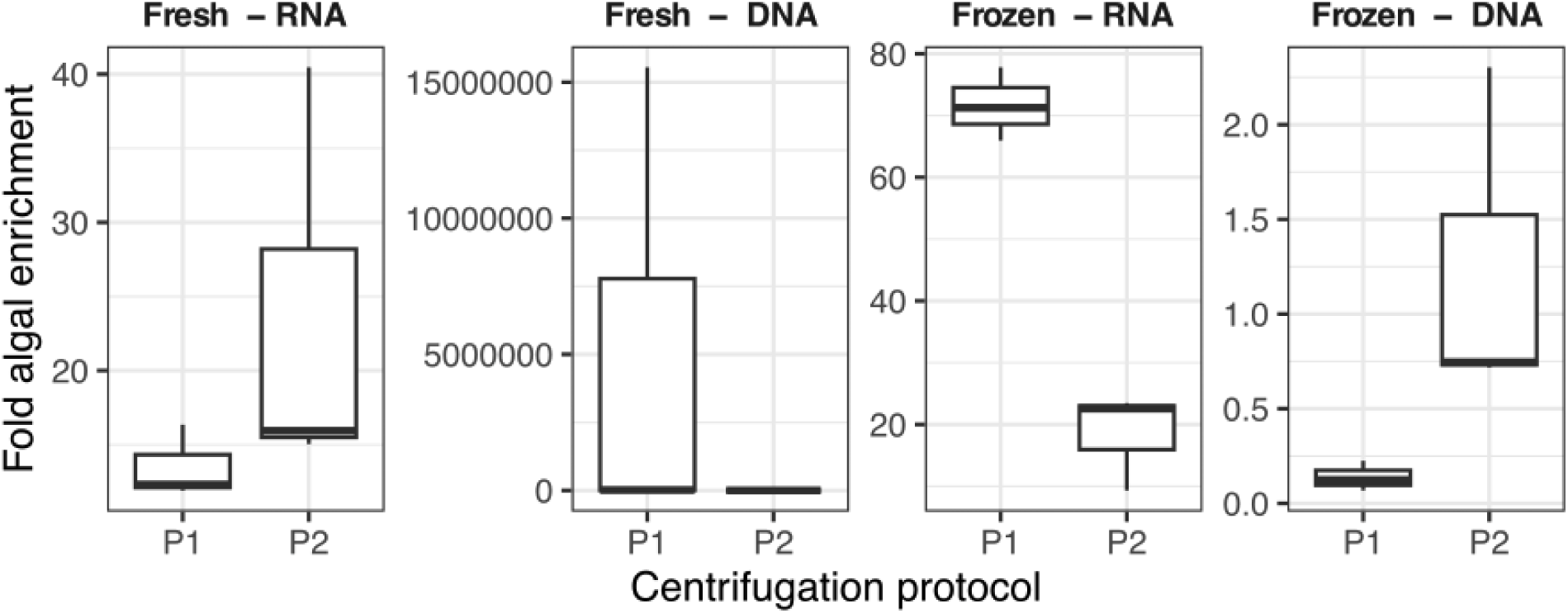
Cell fractionation assessment by amplifying algal- and coral-specific primers using qPCR. Algal enrichment was calculated as the fold increase in algal-to-coral qPCR signal (2^–ΔCt) in fractionated samples relative to unfractionated controls.

**Figure S3.**
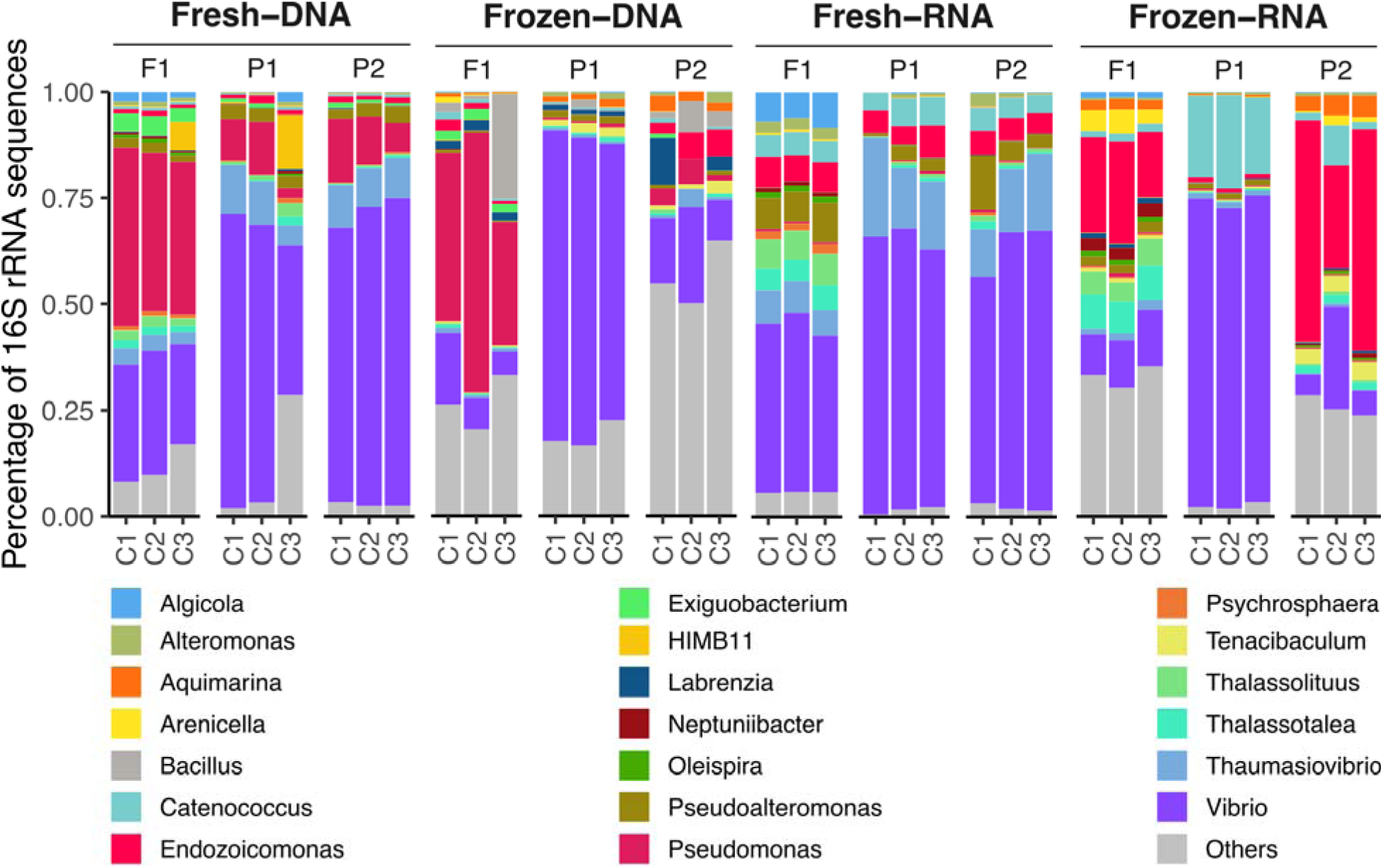
Relative abundance of top bacterial genera across samples. Relative abundance of bacterial genera of unfractionated (F1) and Symbiodiniacea-enriched fractions (F2), across centrifugation treatments (P1 and P2), fresh and frozen samples, and DNA and RNA templates. All other genera were clustered into the Others category.

**Figure S4.**
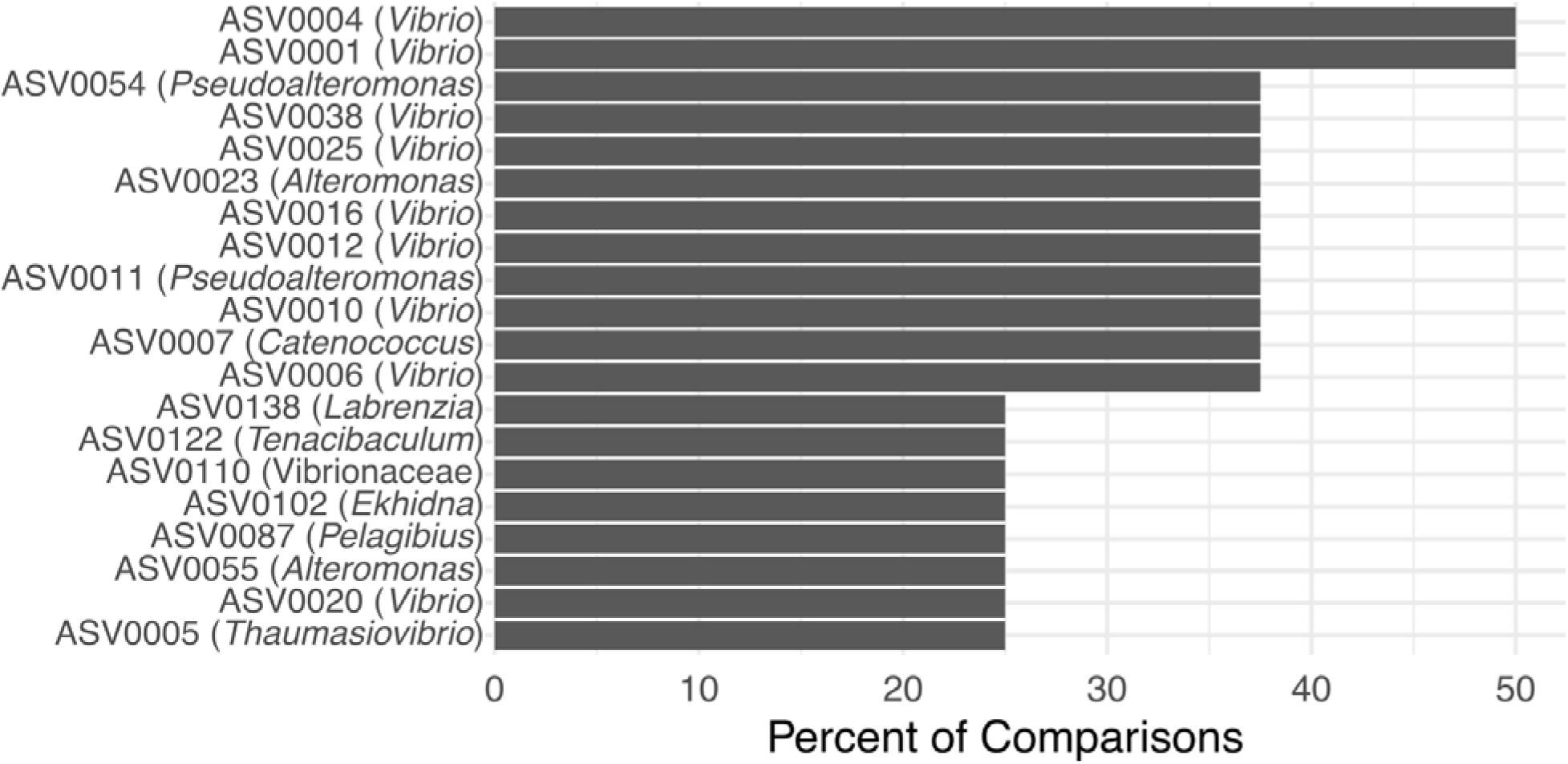
Top 20 bacterial ASVs most frequently enriched in Symbiodiniaceae fractions across treatments. ASV names are displayed along with their assigned genus. Bars represent the percentage of comparisons in which each ASV was identified as significantly more abundant in F2 fractions across the eight treatments, with values sorted in descending order.

**Figure S5.**
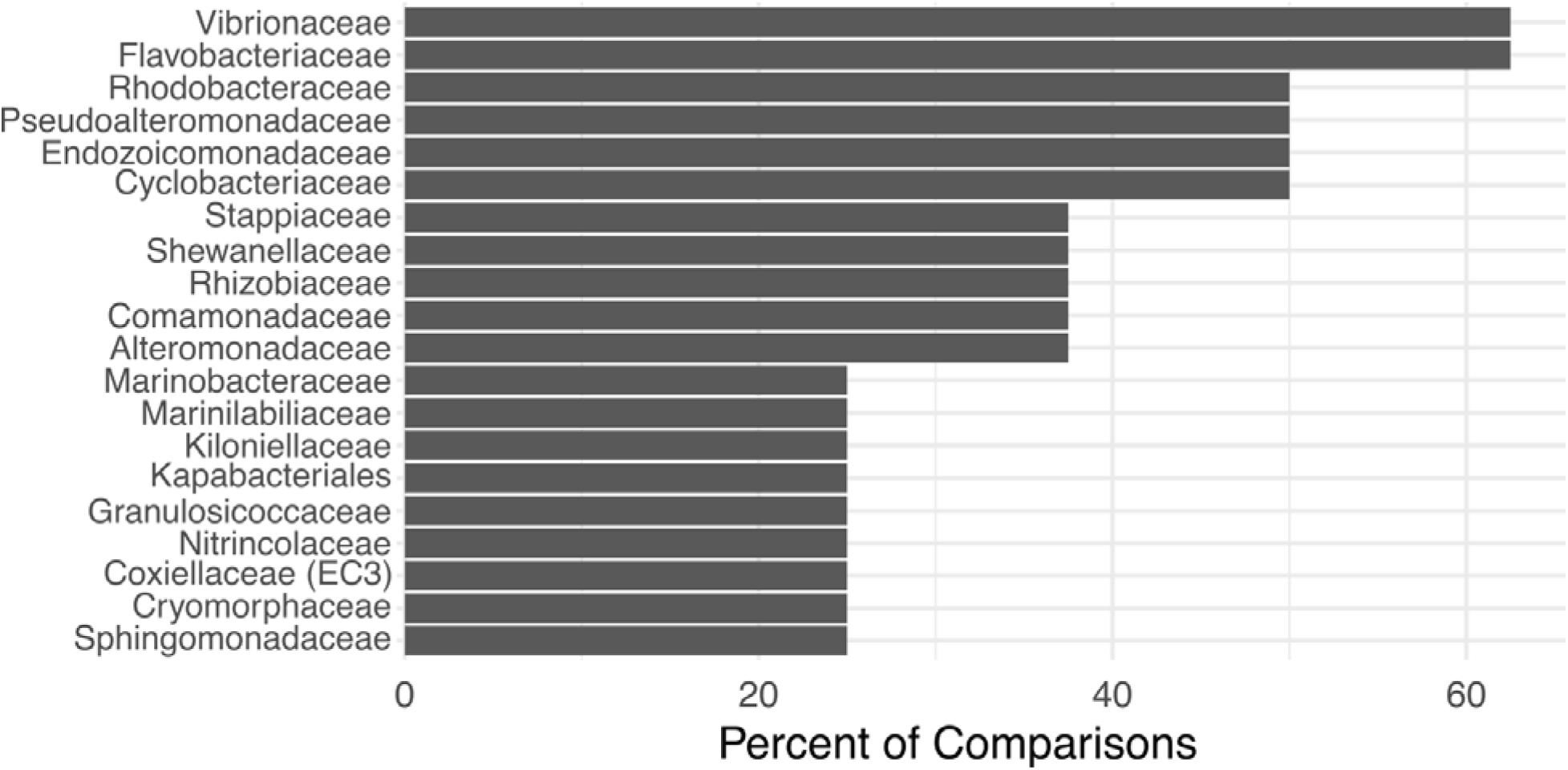
Top 20 bacterial families most frequently enriched in Symbiodiniaceae fractions across treatments. Bars represent the percentage of comparisons in which each family was identified as significantly more abundant in F2 fractions across the eight treatments, with values sorted in descending order. GTR substitution model. The tree was rooted using *Pseudoalteromonas* ASV sequences. Bootstrap support values are shown at selected nodes

**Figure S6.**
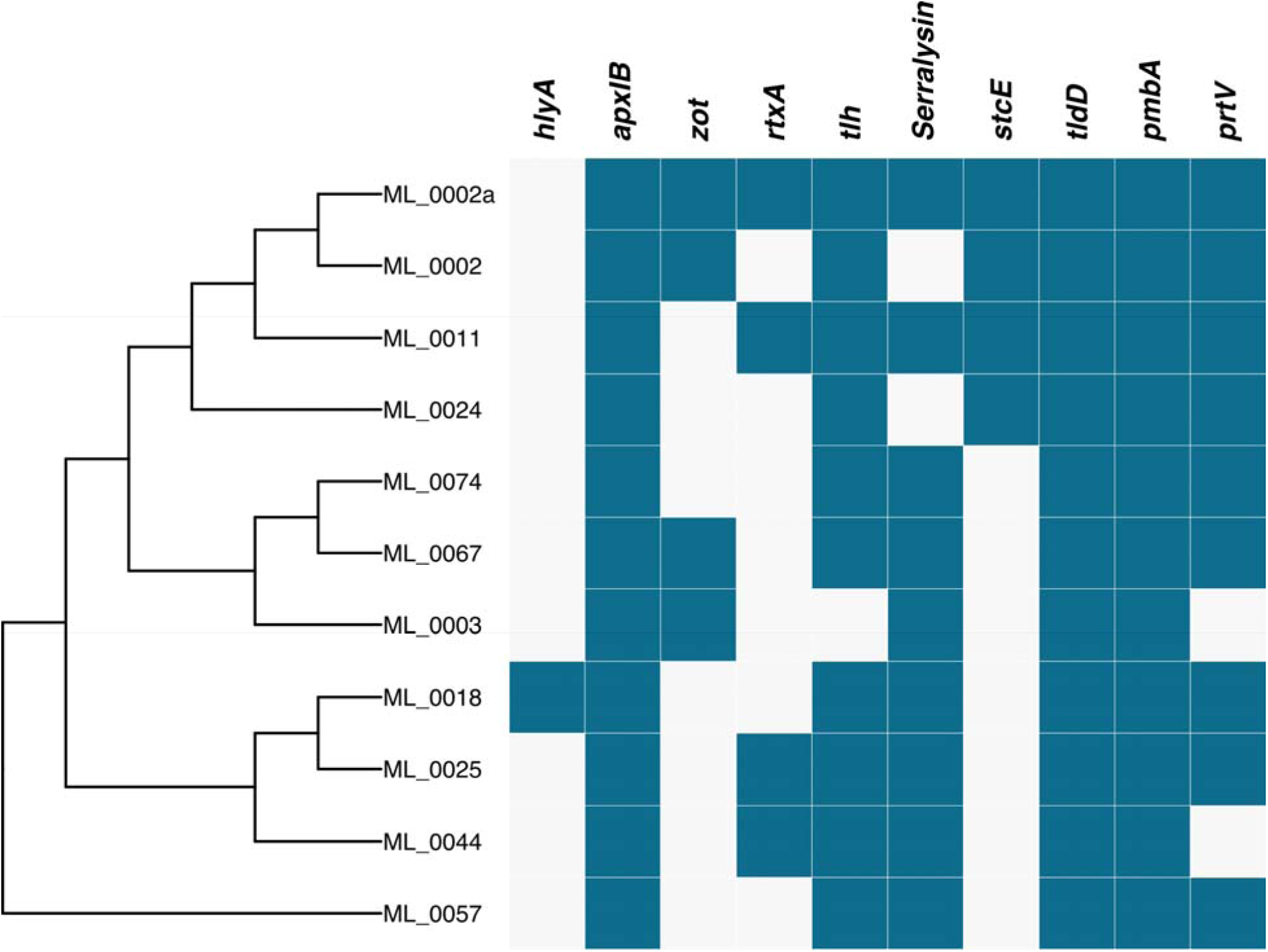
Virulence factors present in the genomes of Symbiodiniaceae-associated *Vibrios*. Phylogenomic tree showcasing the presence or absence of virulence genes in *Vibrios* genomes. Gene presence is represented by a blue square while gene absence is represented by a blank square. Phylogenomic relationships among the 11 Vibrio genomes were inferred using GTDB-Tk.

